# CA1 pyramidal cells organize an episode by segmented and ordered events

**DOI:** 10.1101/565689

**Authors:** Chen Sun, Wannan Yang, Jared Martin, Susumu Tonegawa

**Author notes:** To whom correspondence should be addressed: C.S. or S.T.

## Abstract

A prevailing view is that the brain represents episodic experience as the continuous moment to moment changes in the experience. Whether the brain also represents the same experience as a sequence of discretely segmented events, is unknown. Here, we report a hippocampal CA1 “chunking code”, tracking an episode as its discrete event subdivisions (“chunks”) and the sequential relationships between them. The chunking code is unaffected by unpredicted variations within the events, reflecting the code’s flexible nature by being organized around events as abstract units. The chunking code changes accordingly when relationships between events are disrupted or modified. The discrete chunking code and continuous spatial code are represented in the same cells, but in an orthogonal manner, and can be independently perturbed. Optogenetic inactivation of MEC inputs to CA1 disrupts the chunking but not spatial code. The chunking code may be fundamental for representing an episode, alongside codes tracking continuous changes.

## Main text

How is an episode represented in the brain? It has long been thought that the hippocampus (*1*) encodes the space, objects, and time (*2-4*) in daily episodes. Early studies revealed hippocampal cells that code for variations in space (*5*). Subsequently, hippocampal CA1 cells have been found to be tuned to variations of non-spatial modalities such as passing time (*6, 7*) and variations in sensory stimuli (*6-11*). As a result, a unified view of the hippocampus has emerged as a sequence generator that encodes an episode by tracking its moment-to-moment continuous variations in variables such as space, passing time, and sensory stimuli.

In parallel, psychologists and others, have theorized that episodes are fundamentally subdivided by the brain into events or chunks (*2, 4, 8-11*). Indeed, in real life, rather than remembering all the moment-to-moment continuous variations in their spatio-temporal domain, a typical episode, for instance attending a dinner party, is remembered in terms of segmented events: being led to their designated table, ordering from the menu, waiting for the food, enjoying the jazz band, doing this, and then doing that, etc. An every-day analogy to understand the importance of chunking is to consider the use of folders to organize all the raw documents and data that resides in the computer. Without the widespread use of folders to subdivide computer data into chunks, the organization of data would be an unworkable mess. In other neural systems, visual scenes are segmented by the brain at the highest processing level into discrete objects (*12, 13*), and motor sequences produced by the brain rely on discrete motor sub-programs (*14, 15*). These examples reflect discretization as a general organizational principle of both the computer, and perhaps the brain.

Whether episodic experience, too, is fundamentally segmented by the brain into chunks is not known. The present study aims to investigate whether the hippocampal CA1 subfield carries a code organized around events as fundamental units of episodic experience, supported by the sequentially ordered relationships between these events. Although episodic experience is behaviorally continuous, a chunking process in the brain should allow it to flexibly and efficiently code wide variations in the episodic experiences by organizing around meaningful events, above every moment’s detail. Such an event tracking code could be one of the fundamental organizing principles of episodic experience by the brain.

### Task design to study segmentation of episodic experience into events and relations between them: revealing the “chunking code”

An ordinary episodic experience contains spatial and object variations (Fig. 1a top, middle). We designed our behavior task to be a ‘skeletal’ version of ordinary episodic experience: a sequence of events stripped of differences in spatial and sensory variations to minimize their influence on episodic representations (Fig. 1a, bottom). In our task episode, mice repeatedly ran through a square maze with four laps per trial (Fig. 1d) driven by delivery of a reward at the onset of lap 1 only. These mice visited the reward box after every lap regardless whether a reward was delivered or not (Extended Data Fig. 1a) before starting the next lap. There were two experimental purposes for this task design. *First*, despite the task experience being continuously run, we aimed to see whether a neural code could be found that organized this continuous episode (Fig. 1a top) based on what may be regarded as subdivisions of the task experience: the lap events (Fig. 1a bottom).

**Figure 1:**
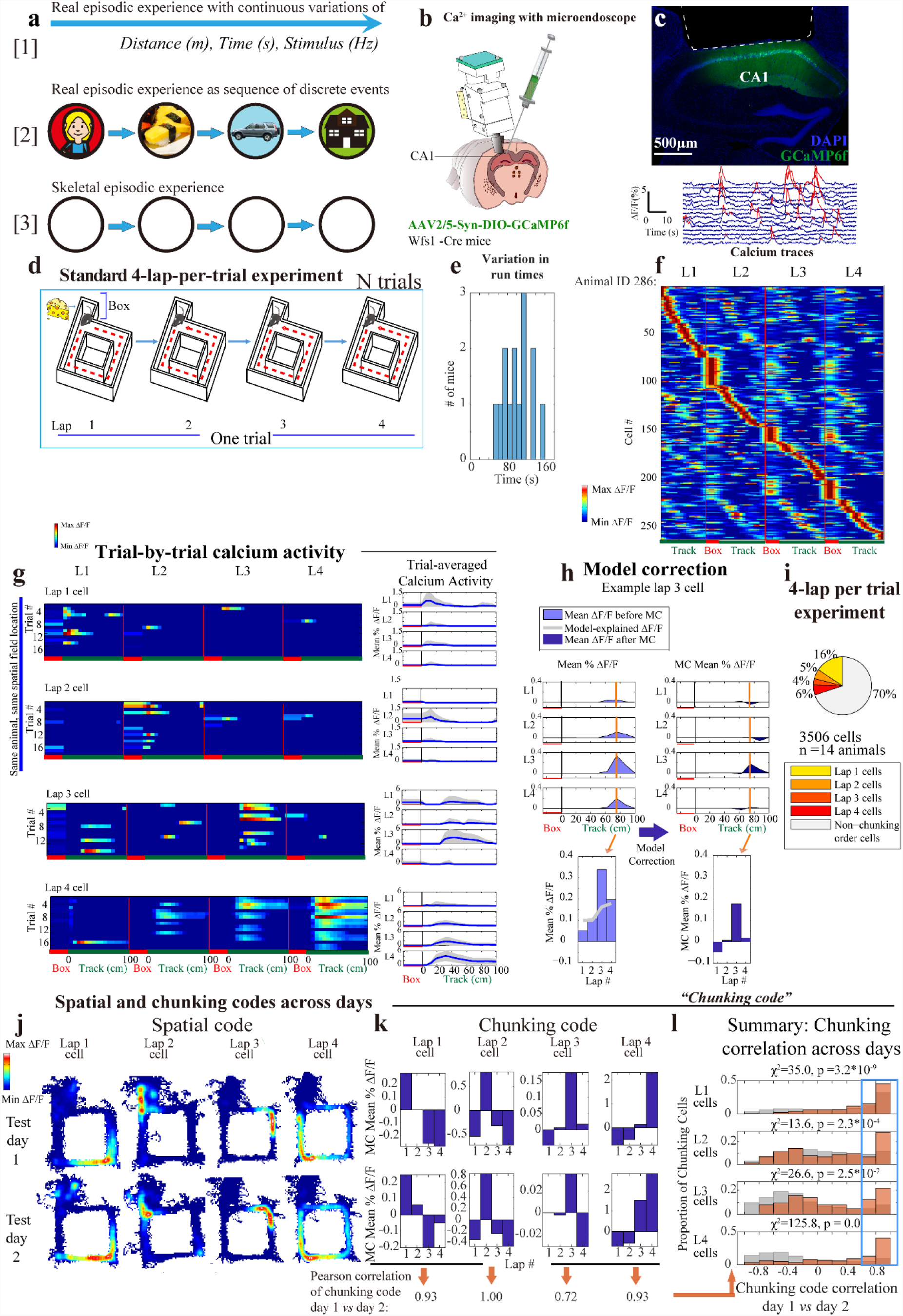
Experimental design to study segmentation of episodic experience. a) Illustration of episodic experience as (**top**): sequence of continuous, moment to moment variations (**middle)**: sequence of discretely segmented events. (**bottom**): Skeletal task experience stripped of sensory and spatial differences, still discretely segmented. b) Implantation of microendoscope into dCA1 of Wfs1-Cre mice with AAV2/5-Syn-DIO-GCaMP6f virus injected in dCA1. c) **Top**: Coronal section of hippocampus showing area of cortex aspiration (white line) and Wfs1+ cells labelled (green). **Bottom**: ΔF/F calcium traces of Wfs1+ (pyramidal) cells in CA1, where (red) denotes significant calcium transients identified. d) During the standard 4-lap-per-trial experiment reward was delivered to the animal at the beginning of lap 1 in the reward box, once every 4 such laps. e) Mean run time among trials (n = 14 animals); f) CA1 calcium activity sorted by spatial position and lap number, normalized and Gaussian smoothed (σ = 25cm) (263 cells from example animal). **Red** label indicates reward box spatial bin, and **green** label indicates the 100 cm long maze track. Reward box activity during lap 1 (reward eating period) was excluded. g) Trial-by-trial calcium activity of representative neurons preferring lap 1, 2, 3, and 4, with spatially binned calcium activity along the track (reward box, plus 16 spatial bins along 100cm track). **Left** panel: trial-by-trial calcium activity; **Right** panel: trial averaged calcium activity (mean ±SD). Standard deviation was cut off at 0 because negative activity does not exist). h) **Left**: representative neuron with raw calcium activity strength sorted by lap number (**Light blue**), and plotted with calcium rate explained by the speed and head orientation fitted linear model (**Grey** trace, See Methods). **Right**: Lap specific remaining calcium rate after the linear model was subtracted, resulting in ‘model corrected’, or MC, calcium activity. **Orange:** peak spatial bin to examine lap modulation of calcium activity i) Summary statistics: Percentage of chunking cells in the whole CA1 pyramidal population that were tuned to lap 1, 2, 3, or 4, in the standard 4-lap experiment (n = 14 animals). j – k) Representative lap 1, 2, 3, and 4 neurons matched across 2 consecutive test days showing spatial code (j) and chunking code (k), as measured by MC calcium activity. For each example cell, Pearson correlations between its chunking codes across days were computed (k, bottom). l) Summary data: the chunking code for individual chunking cells, Pearson correlated across days, plotted separately for lap 1, 2, 3 and 4 cell populations. (**Orange**): Chunking activity of each cell on day 1 correlated with its own chunking activity on day 2. (**Grey**): Chunking activity of each cell on day 1 correlated with chunking activity of arbitrary cells (i.e. shuffled cell identities) from day 2. The proportion of cells with highly correlated (i.e. highly preserved) chunking code across days (cells with Pearson’s *r* > 0.6 threshold: i.e. within the **Blue** box) was significantly greater compared to shuffles: χ^2^ and p values shown in the figure (622 cells, n = 8 animals). See Methods for detailed calculations.

By design, such a neural code would not be tracking different sensory cues since sensory information was identical between the laps, unlike tasks used in other studies (*5-7, 16-26*) in which different events reflect different, constant sensory presentations of visual cues, tones, or odors (Illustration Fig. 1a middle). Rather differently, in our task design, the lap events would reflect a more abstract entity, as we will show. *Second*, having established events as fundamental organizing units of episodic experience, in our particular task the lap events would then reflect a further special property within the experience. Since each of the four lap events was materially identical to one another, if hippocampal CA1 neurons differentially code lap 1, lap 2, lap 3, and lap 4, it would be a code that reflects the abstract sequential relationships between otherwise identical events. To illustrate, representing lap *n* reflects the pure, iterative relationship to the previous lap *n-1* and the subsequent lap *n+1*. Such a representation of abstract sequential relationships would reveal an organizational scheme of episodic experience based on these events.

A virus expressing the calcium indicator GCaMP6f (AAV2/5-Syn-DIO-GCaMP6f) (*27*) was injected into dorsal CA1 (dCA1) of the hippocampus in Wfs1 (Wolframin-1) promoter-driven Cre transgenic mice (Fig. 1b) (*28, 29*). A microendoscope was implanted above dCA1 (*30*) to enable long-term calcium imaging in freely moving mice (Fig. 1b, c). We recorded calcium activity and characterized the spatial selectivity of CA1 neurons (Extended Data Fig. 1c) as mice ran the square maze task (Fig. 1d). During testing, animals completed 15-20 trials in succession. On average, test mice took 98 seconds to complete one trial (Fig 1e). For each neuron during each of the four laps, we calculated its average calcium activity strength during moving periods (> 4 cm/s) to identify differences in calcium dynamics that were related to the lap number (Fig. 1g). Some neurons were found to be most active during reward consumption (lap 1) in the reward box (Extended Data Fig. 1d: representative neurons); these cells were excluded from further analysis because they were active in direct response to the reward (Methods). In general, neurons that were active in the start box during non-rewarded laps, or in the maze, were active at the same location on every lap (Extended Data Fig. 1b, e), but showed a preference for a specific lap (Fig. 1g) with stronger activity than in other laps. We calculated and plotted the trial-by-trial calcium activities of representative neurons for each of the four laps. Representative neurons preferred a particular lap, reliably across individual trials (Fig. 1g, Extended Data Fig. 2). Two representative cells that were observed together in the same animal during the same experiment were active in the same spatial location on the track, but were preferentially and consistently active on different laps (Fig. 1g: lap 1 vs lap 2 cell from animal 197; Extended Data Fig. 2: lap 2 vs lap 4 cell from animal 285).

**Figure 2:**
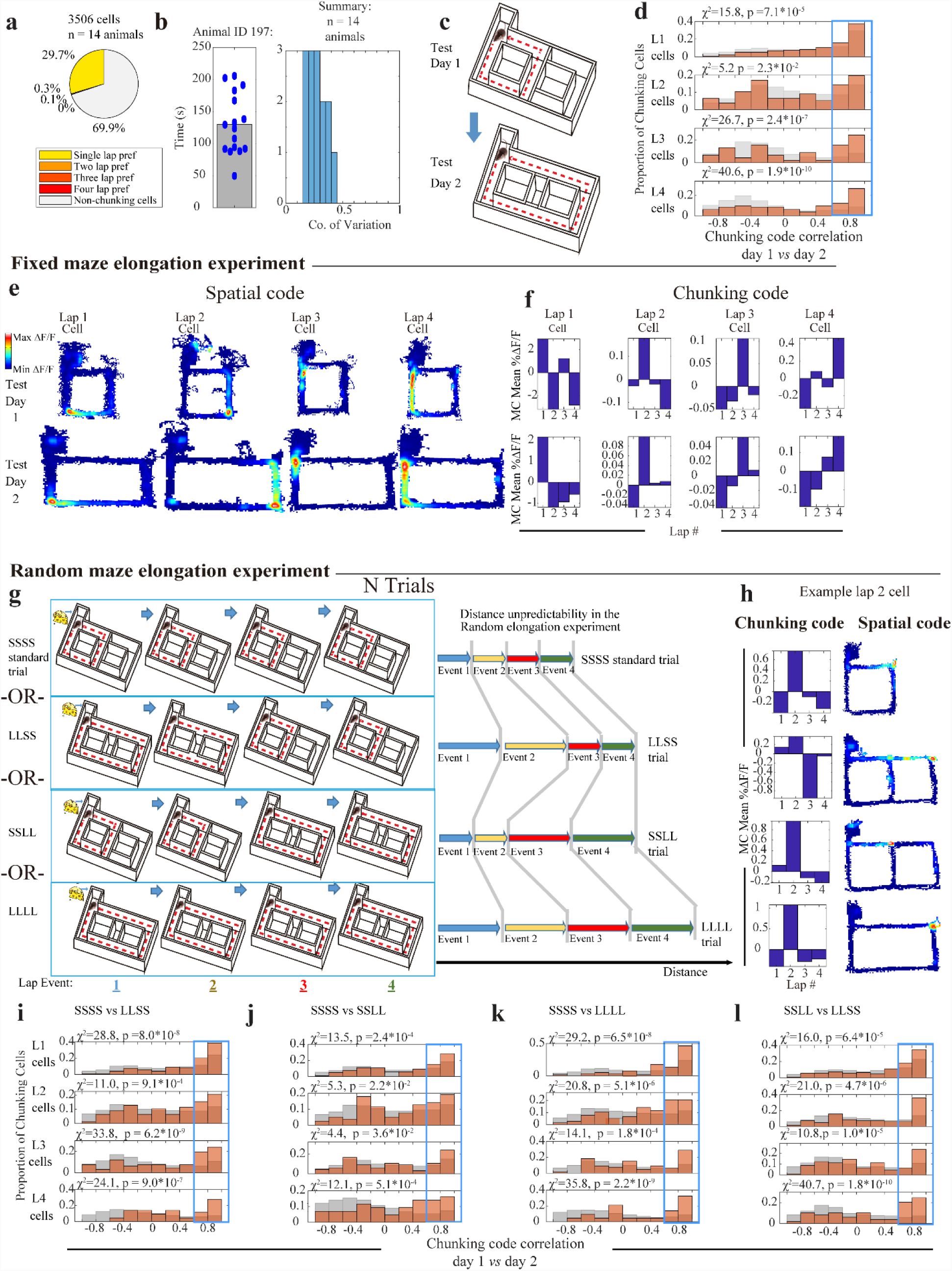
The chunking code is unaffected by variations within events. a) Most chunking cells had statistically enhanced activity on only one lap. b) **Left**: Distribution of running time across trials of animal 197 during 18 trials; **Right**: Coefficient of variation 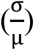 for run time among trials, (n = 14 animals). c) Fixed maze elongation experiment: 4-lap-per-trial task on: Day 1: the standard maze and Day 2: the elongated maze. d) Chunking code correlations across standard and elongated maze sessions (448 cells, n = 6 mice). See Fig. 1(l) for description and methods. e– f) Representative lap 1, 2, 3, and 4 neurons matched across standard and elongated sessions. g) **Left:** Task schedule design for the random maze elongation experiment, where laps were randomly elongated every 2 laps. **Right:** Illustration of the random maze elongation experiment with consistent 4-laps per reward despite variability within the lap events. S denotes a “short” lap. L denotes a “long” lap. h) Example lap 2 cell: its chunking code and spatial code during SSSS, LLSS, SSLL, and LLLL trials (top to bottom respectively). i—l) Chunking code correlations: (i) standard 4-lap trials (SSSS) vs LLSS trials, (j) SSSS vs SSLL trials, (k) SSSS vs LLLL trials, or (l) SSLL vs LLSS trials. (331 cells, n = 6 mice). N.S. denotes ‘non-significant’. See Fig. 1(l) for description and methods.

We calculated the calcium activity across the four laps within spatial bins that tiled the maze (Methods). Since CA1 activity is sensitive to a variety of behavioral variables including spatial location (*5*), running speed (*31, 32*), and head direction (*32, 33*) (Extended Data Fig. 3a-b), we fitted the activity of each neuron to a linear model incorporating the animal’s spatial location, head direction, and running speed (Methods) to account for these variables. We then asked whether these modelled variables were enough to account for the systematic variation in CA1 activity. Thus, we calculated the remaining calcium activity across four laps that was not accounted for by the model and referred to this activity to as ‘model corrected’, or MC, calcium activity (Fig. 1h). Thirty percent (1055/3506 cells, n = 14 mice) of CA1 pyramidal cells had peak, lap-specific MC activity that was significantly different (outside the 95^th^ confidence interval) compared to shuffles. These cells are henceforth called ‘chunking cells’ because their activity is modulated by the lap events (i.e. the chunks) that make up the episode.

**Figure 3:**
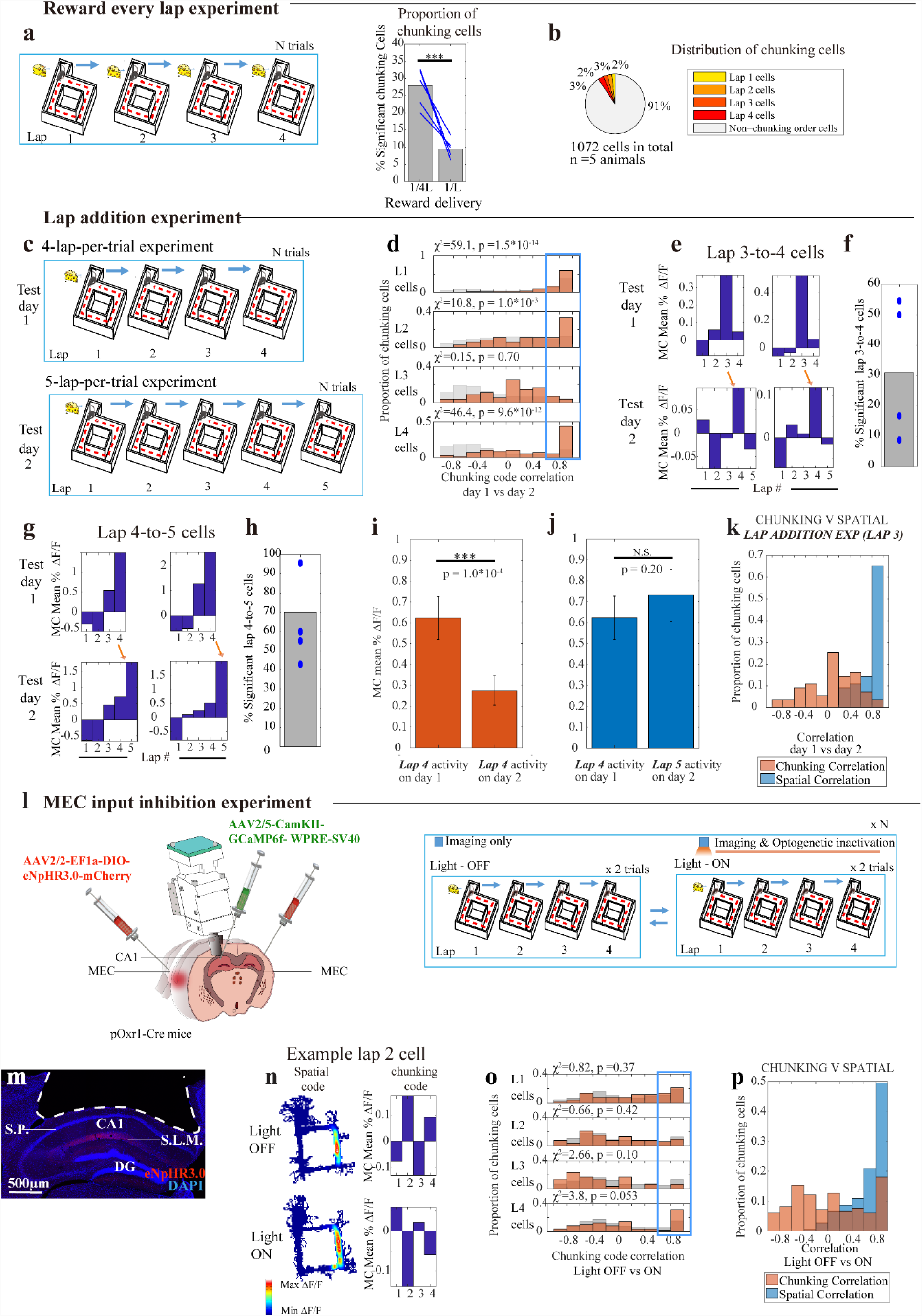
Chunking code tracks the sequential relationship between events and relies on entorhinal input. a) **Left:** Task schedule: reward was given to the animal every lap. **Right:** The percentage of significant chunking cells was significantly reduced in the reward-every lap task, compared with the same animals running the standard 4-lap-per-trial task (**Blue** lines: 5 mice, *χ*^2^ =128.7, p = 0.0). b) Summary statistics: Percentage of chunking cells in the whole CA1 pyramidal population that were tuned to lap 1, 2, 3, or 4, during the reward every lap experiment. c) Lap addition experiment: Day 1: 4-lap-per-trial experiment and Day 2: 5-lap-per-trial experiment. d) Chunking code correlations across the 4-lap and 5-lap experiment sessions (382 cells, n = 4 mice). See Fig. 1(l) for description and methods. e) Two representative neurons matched across 4-lap and 5-lap experiment sessions that transformed from lap 3 to lap 4 preference. f) Percentage of cells that transformed from lap 3 to lap 4 preference (**Blue** marks: 4 mice). g - j) (g) Two representative neurons matched across 4-lap and 5-lap experiment sessions that transformed from lap 4 to lap 5 preference. (h) Percentage of cells that transformed from lap 4 to lap 5 preference (**Blue** marks: 4 mice). (i) MC activity of these cells from (h) during lap 4 on day 1 is significantly decreased during the same lap on day 2. (j) MC activity of these cells from (h) during lap 4 on day 1 is not statistically different from MC activity during lap 5 on day 2 (Mean ±SEM, Paired student t-test conducted). k) Pearson correlation of lap 3 chunking cells’ **Blue**: spatial code and **Orange**: chunking code across days during the lap addition experiment. See Fig. 1(l) for description and methods. l) **Left:** Viral injections permitting CA1 imaging and MEC terminal inhibition in CA1, simultaneously **Right:** During the standard 4-lap-per-trial experiment, optogenetics light-On and light-Off conditions alternated every 2 trials, for a total of 32-40 trials. m) Coronal section of hippocampus showing area of cortex aspiration (white line) and MEC inputs labelled (red). *S.L.M. = stratum lacunosum moleculare. S.P. = stratum pyramidale.* n) Example lap 2 cell: chunking code and spatial code during the light-Off vs light-On trials. o) Chunking code correlations across light-On vs light-Off conditions (182 chunking cells, n = 3 mice). p) Pearson correlation of **Blue**: spatial code and **Orange**: chunking code across light-On vs light-Off conditions, for same cells as in (o). *** denotes p < 0.001, * denotes p < 0.05, N.S., not significant.

There were chunking cells that preferred each of the four laps (Fig. 1i). In order to examine the pattern of calcium activity modulated by lap for a given cell, we calculated its lap modulated MC activity in the peak spatial bin (Fig. 1h, bottom; Methods) for all four laps; we call this *sequence of differential activity strengths* across the four laps the ‘chunking code’ (Fig. 1h, bottom). The percentage of chunking cells increased from 17% on the first day to 29% following eight days training on the lap task (see Methods; pre: 176/1008 cells in n=5 mice; post: 335/1168 cells in the same mice; Extended Data Fig. 4; *χ*^2^ =37.9, p = 7.4*10^−10^) showing that the chunking code is learned. We tracked chunking cells across days and saw that their chunking codes were highly correlated even across days (Fig. 1l, Examples: Fig. 1j-k;—Extended Data Fig. 5a for the analogous raw ΔF/F results).

**Figure 4:**
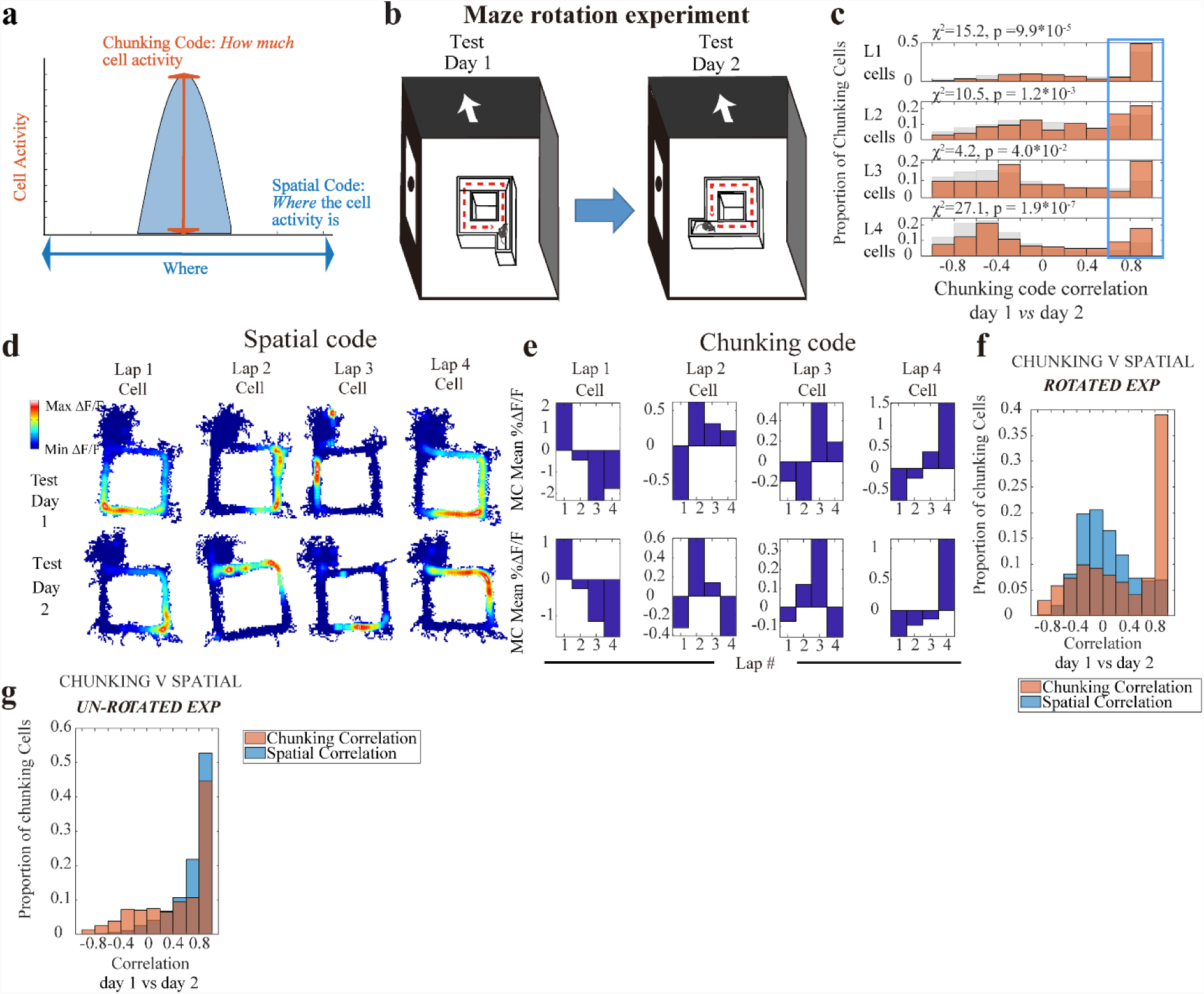
Chunking and spatial codes are jointly represented yet independently manipulatable. a) Illustration: chunking and spatial codes: jointly yet orthogonally represented in the same cells. b) Rotation experiment: Day 1: 4 lap experiment; Day 2: same maze experiment, rotated 90 degrees relative to external cues. c) Chunking code correlations across un-rotated and rotated maze session (692 cells, n = 6 mice). d – e) Representative lap 1, 2, 3, and 4 neurons matched across un-rotated and rotated maze sessions. f) Pearson correlation of spatial code and chunking code where the maze was 90 degree rotated to external cues on Day 2 as (a) (617 cells, n = 5 mice). g) Pearson correlation of spatial code and chunking code across days during the standard 4-lap addition experiment (404 cells in the same 5 mice as (f)).

**Figure 5:**
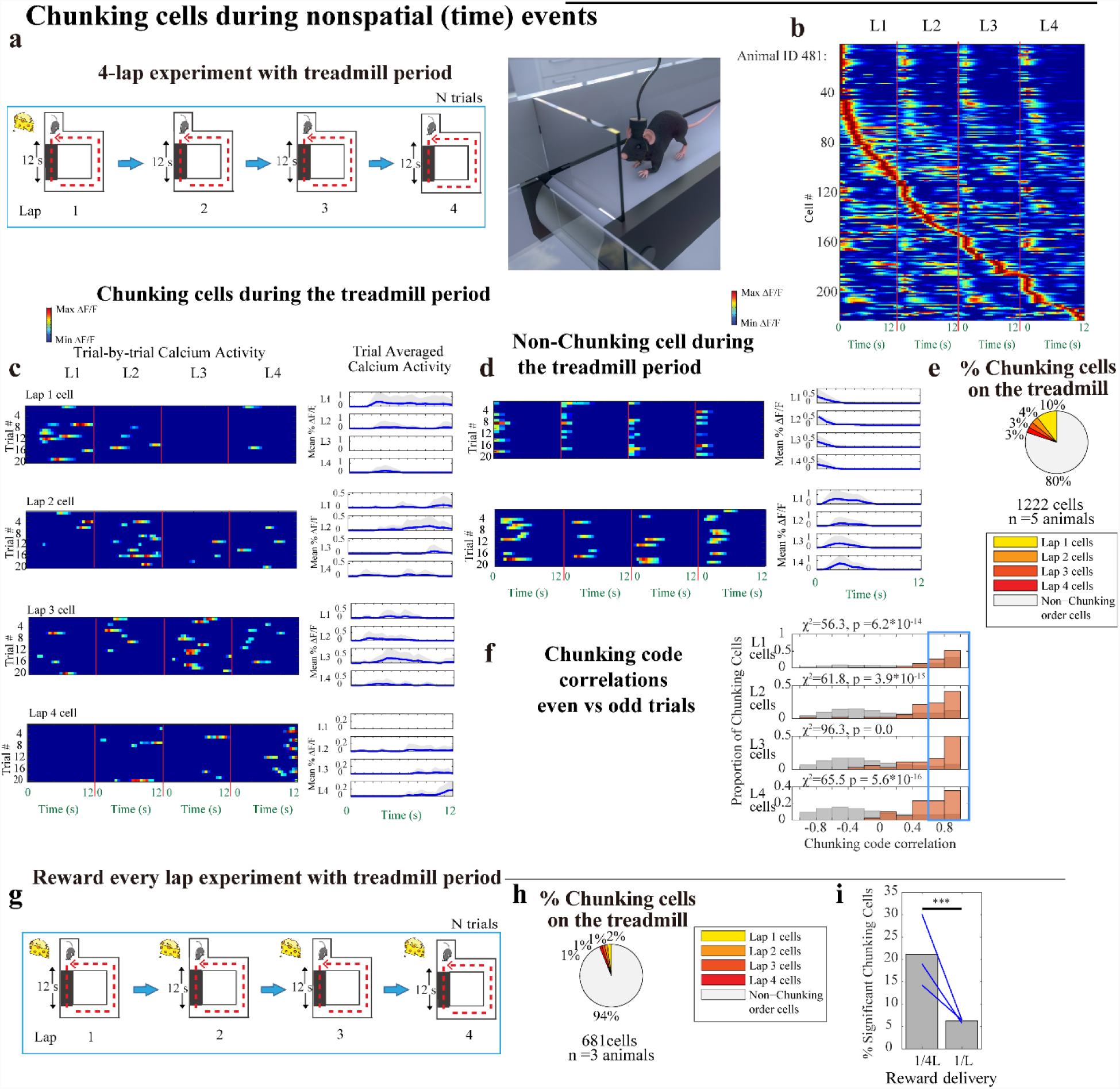
The discrete chunking code is also jointly represented with continuous time code. a) **Left:** 4-lap-per trial experiment with 12s treadmill period on each lap. **Right:** cartoon of mouse running during the treadmill period. The maze and door are not transparent in the task; shown transparent here for illustration of the treadmill below. b) CA1 calcium activity sorted by duration time (s) on the treadmill and lap number, normalized and Gaussian smoothed (σ = 2s) (222 cells from example animal). c) Trial-by-trial calcium activity of representative neurons preferring lap 1, 2, 3, and 4, temporally binned calcium activity during the treadmill period (0.5s bins). **Left** panel: trial-by-trial calcium activity; **Right** panel: trial averaged calcium activity (mean ±SD). Standard deviation was cut off at 0 because negative activity does not exist). d) Trial-by-trial calcium activity of representative neurons that do not have lap preference (ordinary time cells). e) Summary statistics: Percentage of chunking cells in the whole CA1 pyramidal population that were tuned to lap 1, 2, 3, or 4, in the 4-lap treadmill experiment (n = 5 mice) f) Chunking code correlations across between even numbered trials and odd numbered trials (243 cells, n = 5 mice). See Fig. 1(l) for description and methods. g) Task schedule: reward was given to the animal following every lap. Every lap contains a 12s treadmill period. h) Summary statistics: Percentage of chunking cells in the whole CA1 pyramidal population that were tuned to lap 1, 2, 3, or 4, during the reward every lap experiment (with treadmill period). i) The percentage of significant chunking cells was significantly higher during the treadmill period of the 4-lap-per-trial task compared with the same animals during the reward-every lap task (**Blue** lines: 3 mice, χ^2^ =65.0, p = 7.8*10^−16^). *** denotes p < 0.001, * denotes p < 0.05, N.S., not significant.

### The chunking code is unaffected by variations within events

Despite the fact that animal behavior was continuous throughout the task and without experimental breaks, several key results indicated that the chunking code was organized around discrete subdivisions of the episodic experience (i.e. laps). First, although chunking cells were periodically active on each of the 4 laps (Fig. 1f, Extended Data Fig. 1b) the vast majority of them had statistically significant enhanced activity, according to the above criterion, on only one of those laps (Fig. 2a left), indicating relatively sharp lap-specific tuning.

Second, since previous studies (*5-7, 17, 34*) showed hippocampus codes for continuously changing variables, we investigated the chunking code compared to several continuous variable codes more closely. Could chunking cells instead be tracking a particular duration of time since the start of the trial? In fact, episodic or time cells (*6, 7*) require a constant and reliable temporal delay period; otherwise they do not arise (*7*). Because the animals in our task were freely behaving and exhibited unpredictable and variable durations to complete the trials of the task (Fig. 2b), they were unlikely to be time cells. Could chunking cells instead be representing the total distance continually travelled along the course of the 4-lap task since the start of the trial? To test this, we conducted a consecutive two-day experiment in which we elongated the maze in one dimension to twice the usual length on the 2^nd^ day while the animals continued to undergo the standard 4-lap-per-trial task (Fig. 2c, Methods: task specific training). The chunking code was significantly preserved across days (Fig. 2d, Examples: Fig. 2e-f; raw ΔF/F: Extended Data Fig. 5b) despite the spatial distortions of maze length, making it unlikely that chunking cells track the continuous distance traveled.

Finally, real episodic experience has a high degree of variability. To investigate this point further, we next performed a single day experiment with the 4-lap-per-trial task in which the maze was elongated on pseudo-randomly chosen laps of pseudo-randomly chosen trials (Fig. 2g left, Extended Data Fig. 6 for full task schedule, Methods). This maze was largely stripped of predictability in travelled distance but preserved only the 4-discrete lap structure. A total of 26% of CA1 cells (331/1257 cells, n = 6 animals) active in all trial types of this experiment were significant chunking cells. For these chunking cells, their sequence of lap-to-lap pattern of activity strengths (i.e. their chunking code) during the standard (short SSSS) trials was still preserved during each of the pseudo-randomly elongated trial types (Fig. 2i-k; Example cell: Fig. 2h, raw ΔF/F: Extended Data Fig. 5d). For these cells, their chunking codes were even preserved during SSLL trials compared to LLSS trials (Fig. 2l), which were trials that had the same total distance (Fig. 2g right) but differed in their internal segmentation into four lap events. Therefore, the chunking code of this sizeable population of CA1 cells was unperturbed by arbitrary and unpredicted variations within the relevant lap or even variations within neighboring (preceding and ensuing) laps (Fig. 2g right for illustration). These results further indicate that the chunking code treats these lap events as organizing units of the experience.

**Figure 6:**
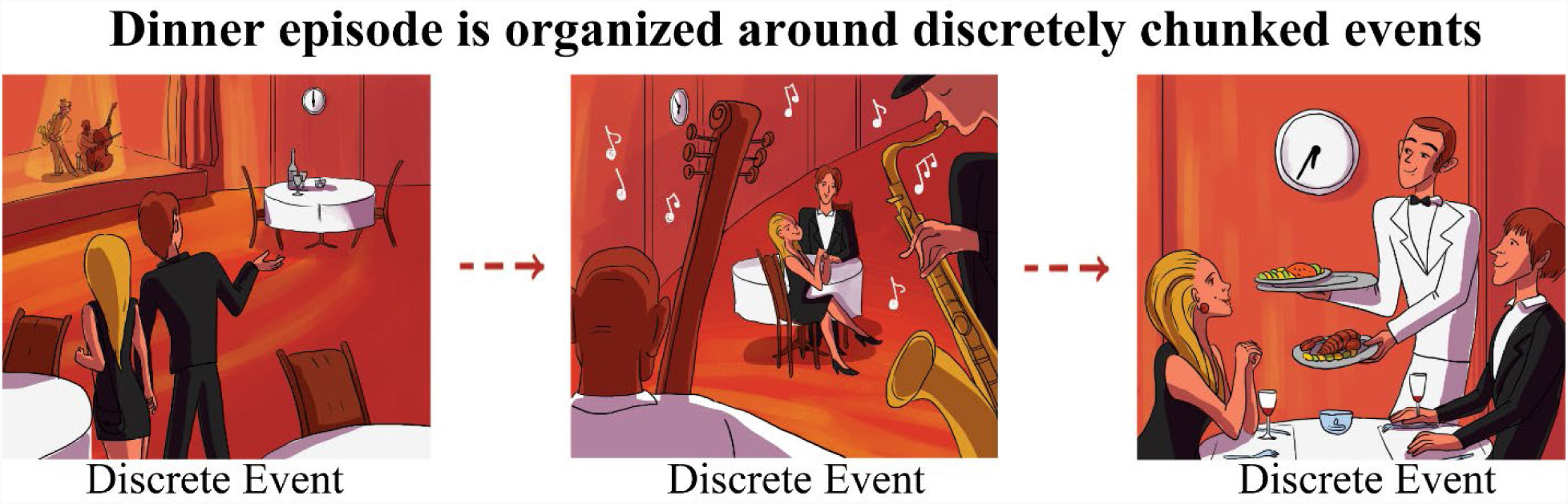
Illustration of a dinner episode. The dinner is organized by the chunking code around discrete events regardless of modality: whether spatial (**left** event: being led the dinner table) or non-spatial (**middle** event: time waiting for the food to arrive, **right** event: enjoying the jazz music) in nature.

It can be seen that the chunking code tracks these lap events in an abstract manner, robust against arbitrary variabilities in physical variables like time (Fig. 2b) or distance (Fig. 2g). We conducted one final examination of the abstract nature of these events. A two day experiment was conducted in which the 4-lap per reward was still preserved on the 2^nd^ day (Extended Data Fig. 7, raw ΔF/F: Extended Data Fig. 5e), whereas the spatial trajectories were altered every two laps (Extended Data Fig. 7, Methods: task specific training) to perturb spatial sequences during the same lap events. Here, a significant proportion of lap 1-4 chunking cells still had preserved chunking code across sessions (Extended Data Fig. 7) despite the animal experiencing differential spatial trajectories. This shows the abstract nature of these “lap events”, tracked by the chunking code.

### The chunking code tracks the sequential relationships between events

Based around these lap events, how is episodic experience then organized? The reliable preservation of lap event-specific activity across trials (Fig. 1g) and across days (Fig. 1l, Examples: Fig. 1j-k) suggests that the chunking code captures sequential relationships between the materially identical lap events. To further test this hypothesis, we conducted two types of experiments.

First, we conducted an experiment in which reward was provided every lap, in order to abolish the sequentially differentiating effect of the once-in-four lap delivery of the reward (Fig. 3a, left). We found that only 9% (101/1072 cells, n= 5 animals, Fig. 3a right, and Fig. 3b) of CA1 cells were significant chunking cells. This was significantly lower than the percentage of chunking cells observed in the standard 4-lap-per-trial experiments in those same animals (28% = 371/1328 cells in the same animals, *χ*^2^ =128.7, p = 0.0).

Second, we conducted a consecutive two-day experiment with the standard 4-lap-per-trial experiment on the first day and added a non-rewarded 5^th^ lap on all the trials of the second day before reward eating (Fig. 3c, Methods: task specific training). We postulated that this subtle modification of the ordering of events may be sufficient to perturb the chunking code. During day 2 of testing, while lap 1 and 2 cells had preserved chunking code despite the additional lap (Fig. 3d; raw ΔF/F: Extended Data Fig. 5c), lap 3 cells were perturbed (Fig. 3d), and a significant proportion of them (17/55 cells = 31%, n = 4 mice, p < 10^−3^ compared to shuffling) shifted to prefer lap 4 (Fig. 3e-f). This was despite the fact that lap 3 and the preceding (lap 2) and succeeding laps (lap 4) were materially identical to each other, as well as across days. Lap 4 cells showed a similarly shifted chunking code to prefer lap 5 (Fig. 3g-h). Although, overall, the pattern of a majority of Lap 4 cell activity across the two days was well correlated across the first 4 laps (Fig. 3d), a proportion of lap 4 cells (42/60 cells = 70%, n = 4 mice, p < 10^−4^ compared to shuffling) were subtly perturbed. These data show a decrease in overall activity strength during lap 4 on day 2 (Fig. 3i) with a concomitant restoration of activity strength during lap 5 (Fig. 3j). Thus, the alterations of the materially identical lap 3 and 4 representations were due to the addition of the 5^th^ lap that changed the sequential structure of the task. Such alterations perturbed lap 3 and 4’s prospective sequential relationships to the end of the trial.

Interestingly, when the sequential organization of events was altered by the addition of this extra lap, the chunking code was correspondingly altered as well, but it did so in a discrete, lap-specific manner rather than gradually through the course of the task experience (Fig 3d). Indeed, whereas lap 1 and 2 representations were well correlated across days, representations during lap 3 were abruptly and discretely altered. This further supports the idea that the chunking code organizes episodic experience around discrete segments—and even chunking code changes happen to occur along these natural segments of the episode.

Taken together, these data show that the chunking code organizes an episode around events and reflects the abstract sequential relationships between them.

### Chunking and spatial codes are jointly but independently represented in the same cells

What is the relationship between the chunking code for discrete events and the well-known place code for continuously changing space (*5*)? The chunking and spatial codes occur in the same cells, yet are treated in an orthogonal manner: the spatial code manifests as *where* on the spatial track the neuron is active, and the chunking code manifests as *how much* the neuron is active, during each different laps (Fig. 4a), without affecting the spatial tuning during each lap (Fig. 1f, Extended Data Fig. 1b). Within this joint arrangement of the two codes, we further hypothesized that the discrete chunking code could be manipulated independently from the continuous spatial code. When the maze and task were not altered in any way across days, both chunking and spatial representations remained highly correlated (Fig. 4g). In contrast, perturbing the sequential relationships between events by adding a lap (Fig. 3c) altered some chunking representations (Fig. 3d-f, Fig. 3k, orange histogram) but still preserved spatial representations in the same cells (Fig. 3k, blue histogram).

A perturbation of the chunking code can also be seen in a different experiment. Since medial entorhinal cortex (MEC) input into CA1 has been implicated in the sequential organization of episodes (*35-38*), we asked how MEC inhibition might affect the chunking code versus the spatial code. Based on these earlier studies, a virus expressing inhibitory opsin (AAV2/2-EF1a-DIO-eNpHR3.0-mCherry) was injected bilaterally into the MEC sub-region of pOxr1-Cre mice (Fig. 3l left, 3m). In addition, a virus expressing the calcium indicator GCaMP6f (AAV2/5-CamKII-GCaMP6f-WPRE-SV40) was unilaterally injected into dorsal CA1 (dCA1) of the same mice (Fig. 3l left). An opto-endoscope was implanted above dCA1 to enable long-term calcium imaging as well as optogenetic inhibition of the axonal terminals from MEC neurons in dCA1. The mice ran 28-40 trials of the 4-lap task where the trials alternated between inactivation (Light-On) and no inactivation (Light-Off) (Fig. 3l). Inactivation of MEC cell terminals in dCA1 globally altered chunking representations but did not change the spatial representations in the same cells (Fig. 3o-p, Example cells: Fig. 3n, Extended Data Fig. 8). Therefore, the chunking code is different from the spatial code even though they are represented jointly in the same cells.

Can the converse result, spatial code alteration without chunking code alteration, be observed? To investigate this point, we conducted a consecutive two-day experiment in which we rotated the maze by 90 degrees on the 2^nd^ day while the animals continued to undergo the standard 4-lap-per-trial task (Fig. 4b, Methods: task specific training). In this case, the chunking code for lap 1, 2, 3 and 4 cells were all preserved across days (Fig. 4c, Fig. 4f, orange histogram; raw ΔF/F: Extended Data Fig. 5f) despite the spatial rotation of the entire maze. The chunking code preservation was in contrast to a change in spatial field location, relative to the maze, in the same cells (Fig. 4f, blue histogram, Examples: Fig. 4d-e). In this way, the same cell can simultaneously code a different spatial moment within the lap run, and still code the same lap event number.

Thus, these experiments show that the hippocampal CA1 chunking code, manifested as event-modulated activity strengths, is jointly yet orthogonally represented in the same cells as the spatial code, and the two codes can be manipulated independently of one another.

### The discrete chunking code is also jointly represented with the continuous time code

If indeed the brain tracks episodic experience jointly via a discrete chunking code and a continuous spatial code, would it continue to represent episodes in this dual manner even in an episode where the main continuous changes are *not* spatial? To answer this question, we conducted another 4-lap-per-trial experiment using a continuous time code. In this experiment, the first arm of the standard 4-lap-per-trial maze was replaced by a treadmill (Fig. 5a). Animals ran for 12s on a treadmill at 14 cm/s on every lap. Monitoring activity of neurons on a treadmill obviates the necessity of model corrections for running speed and head direction changes (Fig. 1h) because they are nearly constant on the treadmill (Extended Data Fig. 9a-b). The treadmill experiences gave rise to time-modulated cells (*6, 7*) (Fig. 5d) that tiled the 12s experience (Fig. 5b, Extended Data Fig. 9c), just as place cells had previously tiled the length of the maze run (Fig 1f). On the treadmill, 20% of CA1 cells (243/1222 cells, n = 5 animals) had significantly different activity depending on the lap number (i.e. the chunking code) (Fig. 5e, Methods, and examples: Fig. 5c, Extended Data Fig. 9d). It can be checked that the chunking code of these cells during the treadmill period was robustly preserved between trials (Fig. 5f). Note that the percentage of chunking cells was significantly reduced during the control task where reward was given every lap (6% = 42/681 cells, n = 3 animals, Fig. 5g-i). These results show that the simultaneous tracking of episodes in both a discrete manner and a continuous manner is a fundamental organizing principle, regardless whether the episodic experience is primarily spatial or temporal in nature (Fig. 6).

## DISCUSSION

How does the hippocampus encode an episodic experience? The main finding of this study was the identification of a hippocampal code that organizes around discrete subdivisions of the episode (i.e. events) and their sequential relationships to one another—the chunking code. While there has emerged a view that the hippocampus tracks an episode as a sequence generator (*5-7, 16-25, 39, 40*) by tracking the moment to moment continuous variations for both spatial or non-spatial moments, in this study the chunking code tracks the same episodic experience within the same CA1 cells via a different organizing principle: organizing it around discrete chunks of experience and their relationships (Fig. 2). In our task (Fig. 1d) the chunking of experience into events did not require any differential sensory cues (*5-7, 16-25, 39*) to define an event or permit the distinction of these events from one another, nor even require any differential past or future sensory cues (*19,25*). Instead, these events had an abstract nature, unchanged by variations in distance travelled (Fig. 2c-f), time duration (Fig. 2b), changes in spatial trajectories (Extended Data Fig. 7), or even spatial rotation of the entire episode (Fig. 4). Even when the chunking code changed (Fig. 3d), it changed along the segments of the episode—in a discrete, lap-specific manner—rather than gradually through the course of the task experience, showing that a lap event is treated as a discrete, unitary entity by the chunking code.

In the end, organizing an episode based on the discrete chunking code tracking events and their relationships, and a continuous code tracking moment to moment changes during the same episode, coexist in the hippocampus. In fact, the discrete chunking code and continuous code are jointly represented in the same cells for both spatial and non-spatial episodes (Fig. 4-5). This supports the concept that the simultaneous tracking of episodes in both a segmented and continuous manner is a fundamental and general encoding principle, which the hippocampus uses. Nevertheless, the two codes are separate representations: in fact, the chunking code and spatial code can be independently altered without affecting the other (Fig. 3k, 3p, Fig. 4f, 4g). In fact, the two joint codes represent different aspects of the same episodic experience. Indeed, the tracking of immediate experience within an event likely requires a level of detail that would be best served by a continuous (spatial or non-spatial) neural representation. On the other hand, tracking the meaningful episodic events above the moment-to-moment variational details is best served by a flexible and discrete representation (Fig. 6). This study provides experimental evidence for a novel “chunking code” in the hippocampus tracking events and their relationships, which could be a fundamental code by which episodic experience is encoded efficiently and flexibly in the brain.

## Acknowledgements

We thank T. Kitamura, J. Z. Young, D. Roy, M. Wilson, E. Brown, M. Sur, M. Harnett, M. Hasselmo, S. Muralidhar, B. Sun, J. Tao, N. Chen and C. MacDonald for comments; F. Bushard, A. Hamalian, C. Ragion, C. Lovett, D. King, and Ella Maru Studio for technical assistance; L. Brenner for paper preparation, and the members of Tonegawa lab for their support. This work was supported by the RIKEN Center for Brain Science, the Howard Hughes Medical Institute, and the JPB Foundation (to ST).

## Author Contributions

C.S., and S.T. designed the study. C.S., W.Y., and S.T. interpreted the data. C.S. and J.M. conducted the surgeries, behavior experiments, and computational analyses. C.S., W.Y., and S.T. wrote the paper. All authors discussed and commented on the manuscript.

## Data availability

The data and code that support the findings of this study are available from the corresponding authors upon reasonable request.

---

**Extended Data Fig. 1:**
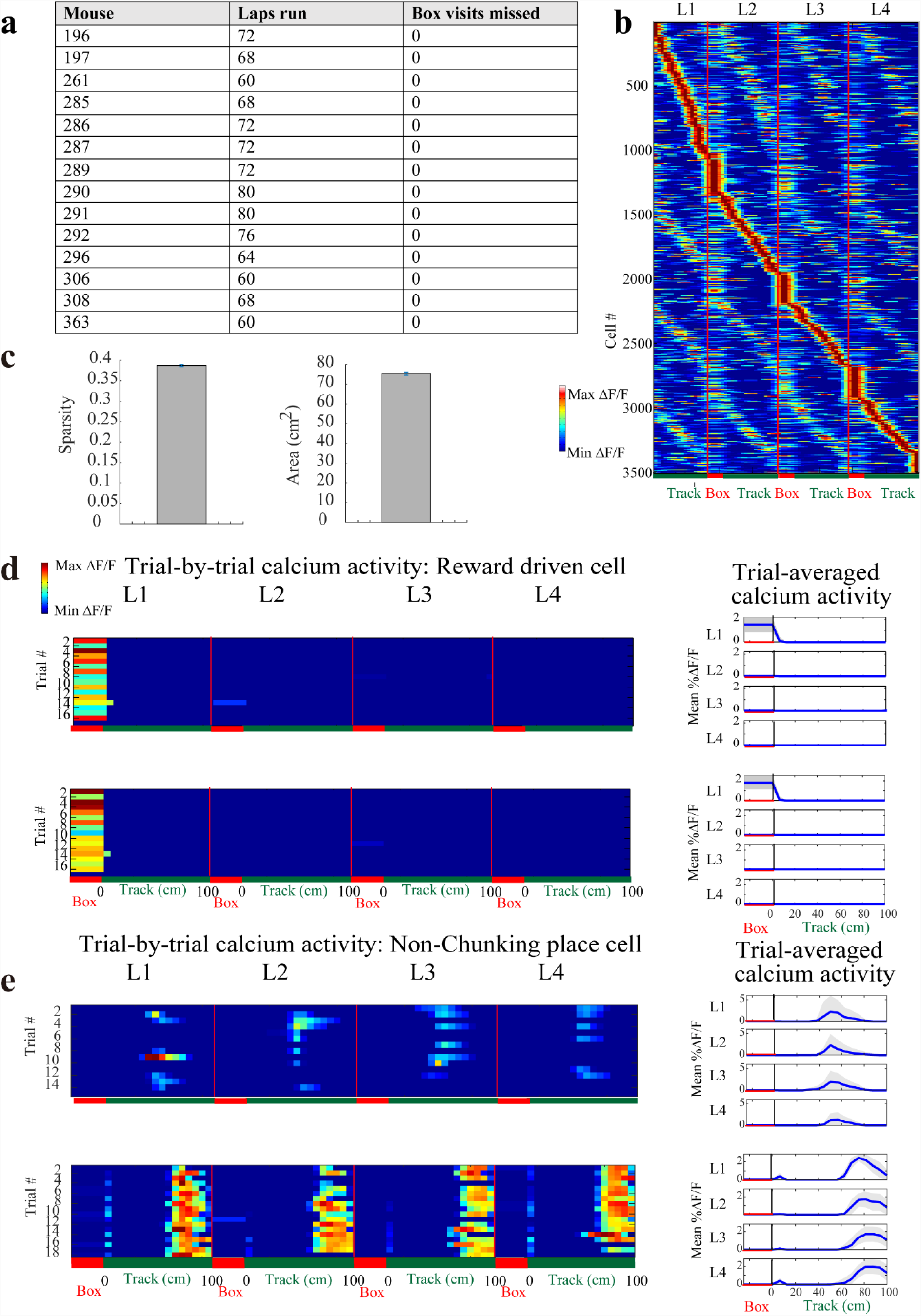
Spatial and Reward properties of CA1 cells on the maze. a) Summary of mice running in a single session of the standard 4-lap-per-trial task. Mice did not miss a visit into the reward box on any run. b) CA1 calcium activity sorted by spatial position and lap number, normalized and Gaussian smoothed (σ = 25cm) calcium activity (3506 cells, n = 14 animals). **Red** label indicates reward box spatial bin, and **green** label indicates the 100 cm long maze track. Reward box activity during lap 1 (reward eating period) was excluded. c) Characterization of mean spatial properties of CA1 cells active in the lap maze: **Left:** sparsity, and **Right:** spatial field size, plotted mean ± SEM; n = 14 mice. In total, 72% (2509/3506) of CA1 cells from 14 animals were significant place cells. d—e) Spatially binned calcium activity along the track (reward box, plus 16 spatial bins along 100cm track) showing (d) 2 representative cells that were active in response to reward, and (e) 2 representative place cells that did not have lap modulated activity. **Left** panel: trial-by-trial activity **Right** panel: trial-averaged activity with mean ± SD. Standard deviation was cut off at 0 because negative activity does not exist.

**Extended Data Fig. 2:**
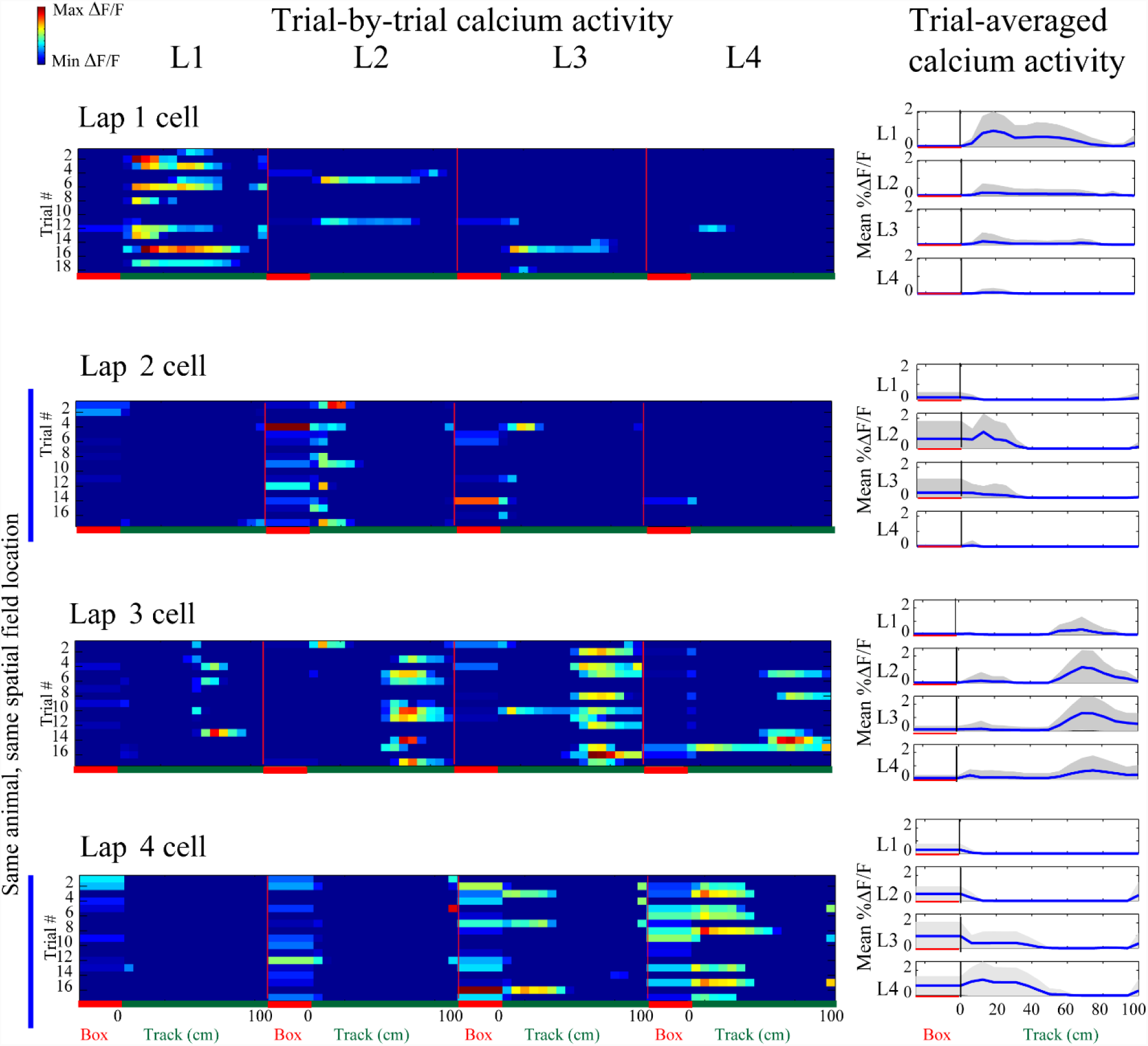
Lap 1-4 cells (more examples) Representative lap 1, 2, 3, and 4 neurons, spatially binned calcium activity along the track (reward box, plus 16 spatial bins along 100cm track). **Left** panel: trial-by-trial activity, **Right** trial averaged activity with mean ± SD. Standard deviation was cut off at 0 because negative activity does not exist.

**Extended Data Fig. 3:**
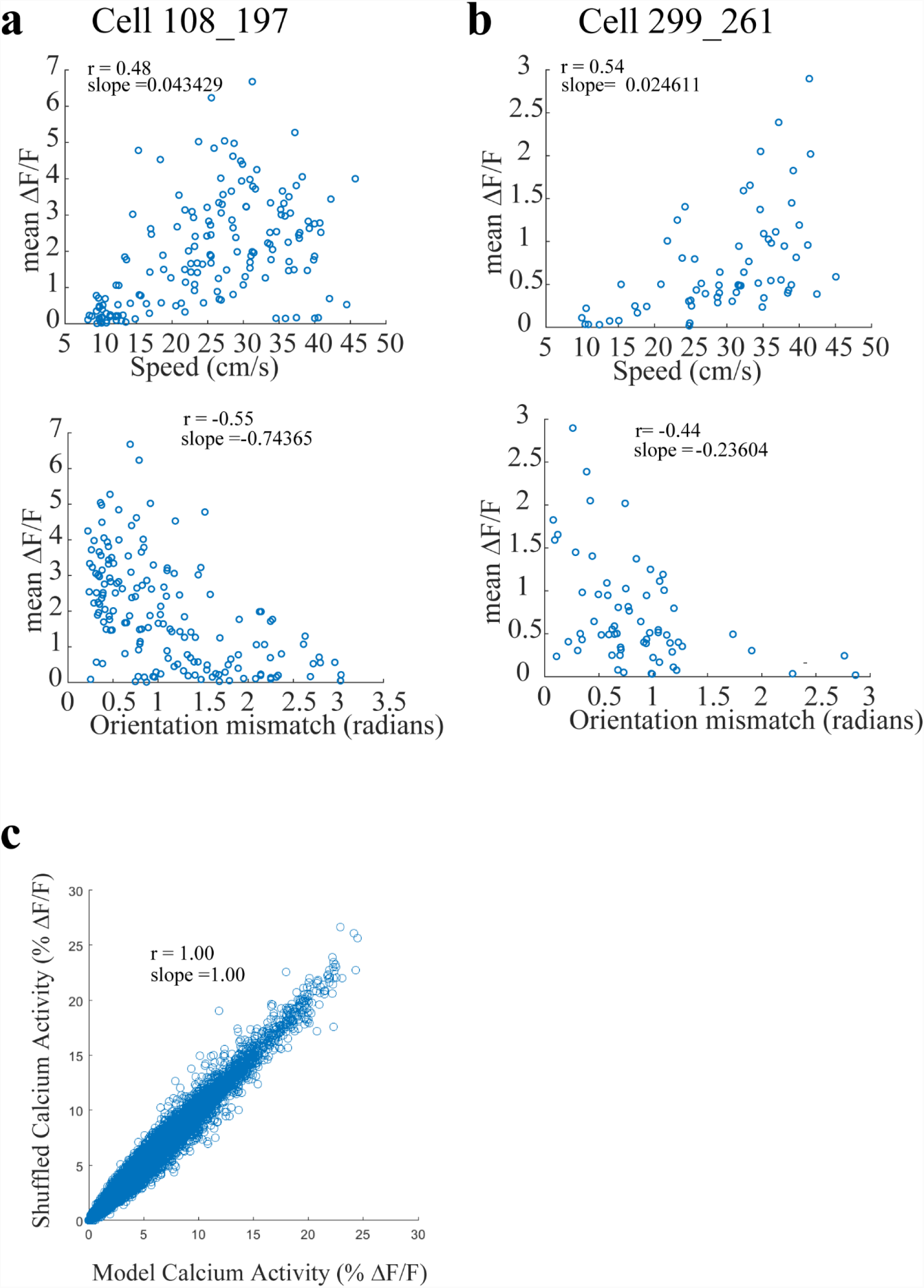
Model correction for Speed and Head direction modulations of CA1 cell activity. a) – b) Two representative cells with calcium activity level plotted against mean animal running speed (top subpanels), and head direction tuning (bottom subpanels). *r* denotes Pearson’s correlation. c) Shuffling procedure preserves the mean calcium activity level as prescribed by the linear model (See Methods) (*r* denotes Pearson’s correlation from 14 animals).

**Extended Data Fig. 4:**
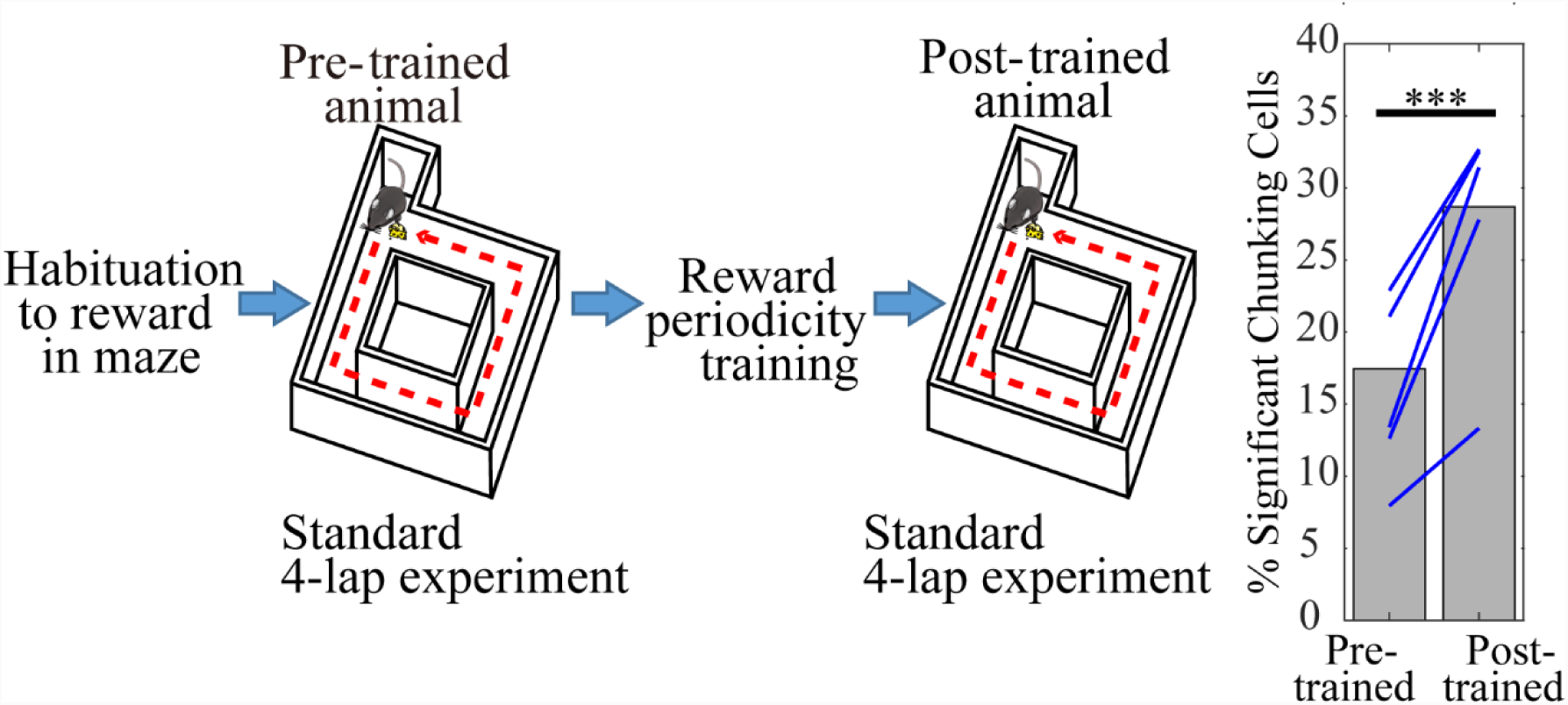
The chunking code is learning dependent. **Left:** Experimental schedule for pre-trained vs post-trained animals on the standard 4-lap-per-trial task. **Right:** The percentage of significant chunking cells was significantly less for pre-trained comparing with post-trained in the same mice (**Blue** lines: 5 mice; *χ*^2^ =37.9, p = 7.4*10^−10^)

**Extended Data Fig. 5:**
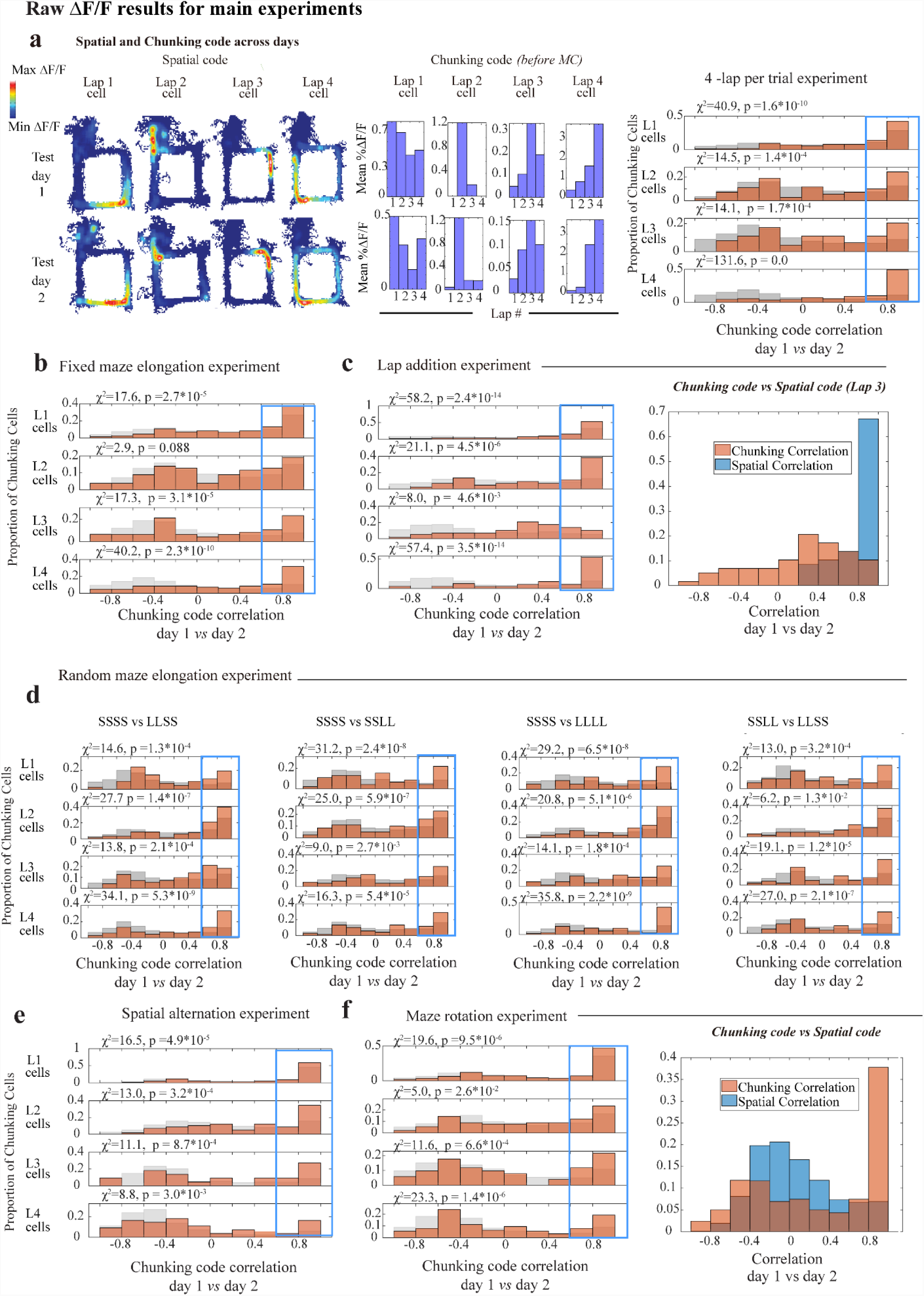
Chunking code preservation across sessions as calculated using raw ΔF/F activity. a) **Left:** The same representative lap 1, 2, 3, and 4 neurons matched across 2 consecutive test days as Fig 1j-k above, measured by raw (i.e. non-model corrected) ΔF/F calcium activity. **Right:** Pearson correlation of chunking code across days, calculated using raw ΔF/F activity. The cells here were the same animals and experimental sessions as Fig. 1l above, plotted separately for lap 1, 2, 3 and 4 cell populations. b) – f) Pearson correlation of chunking code across sessions, calculated using raw ΔF/F activity, for the (b) fixed maze elongation experiment from Fig. 2c-f, (c) lap addition experiment from Fig. 3c-k, (d) random maze elongation experiment from Fig. 2g-k, (e) spatial alternation experiment from Extended Data Fig. 7, and (f) maze rotation experiment from Fig. 4b-f. The cells here were the same animals and experimental sessions as the corresponding plots in the main figures, plotted separately for lap 1, 2, 3 and 4 cell populations.

**Extended Data Fig. 6:**
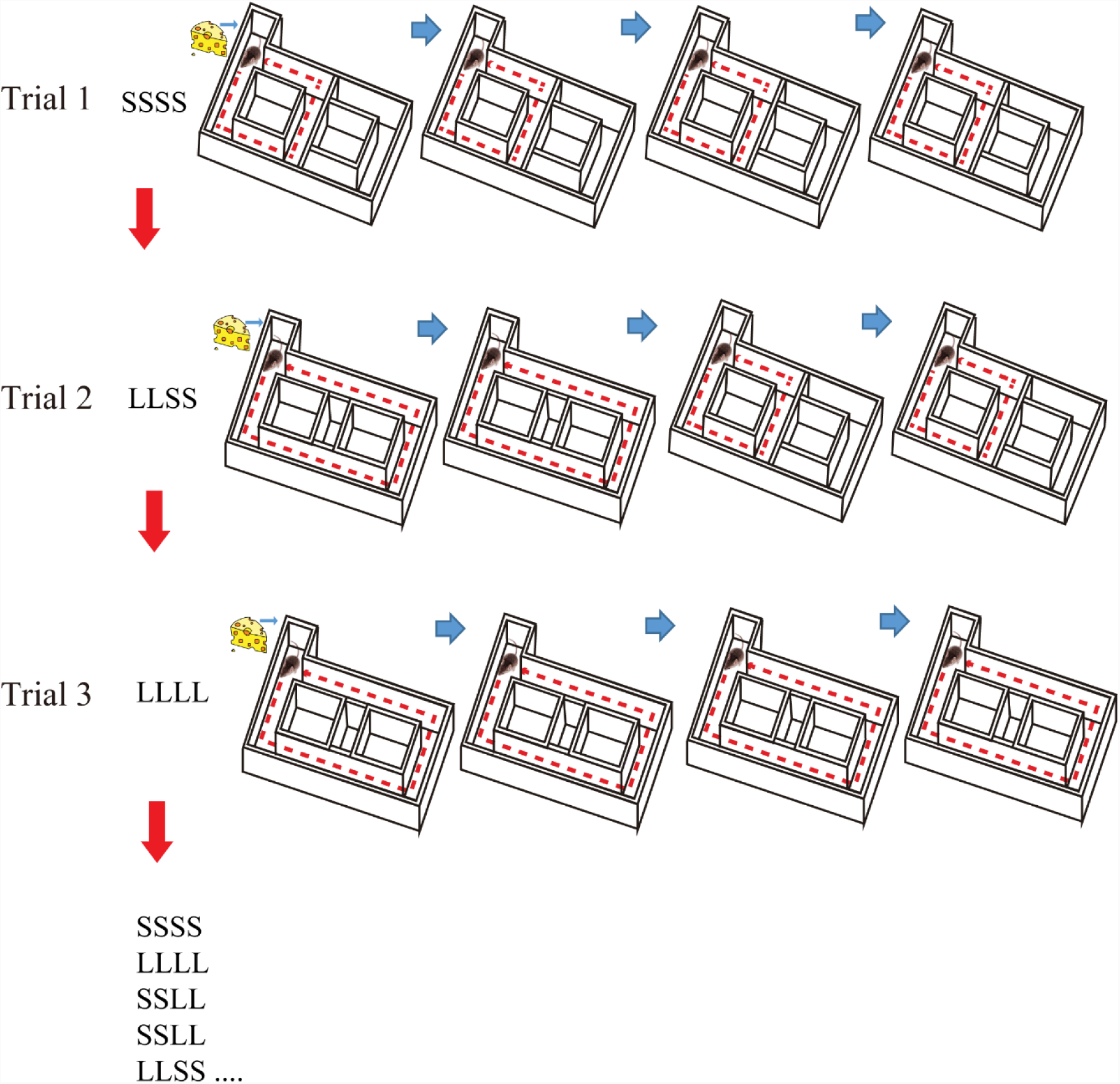
Task schedule for the random maze elongation experiment. The first 3 consecutive trials of the random elongation experiment. Each animal underwent a pseudorandom sequence of 28 consecutive 4-lap trials. Within these 28 trials, 7 trials took place on the standard short maze (SSSS), and 7 trials were of each of the other 3 types SSLL, LLSS, LLLL, where the maze was randomly elongated during L “long” laps. The entire 28 consecutive sequence of trials was: SSSS, LLSS, LLLL, SSSS, SSSS, SSLL, SSLL, LLSS, LLLL, LLSS, LLSS, SSLL, LLLL, LLLL, SSSS, LLLL, LLLL, LLSS, LLLL, LLSS, SSSS, LLSS, SSLL, SSSS, SSSS, SSLL, SSLL, SSLL in this order.

S denotes a “short” lap and L denotes a “long” lap.

**Extended Data Fig. 7:**
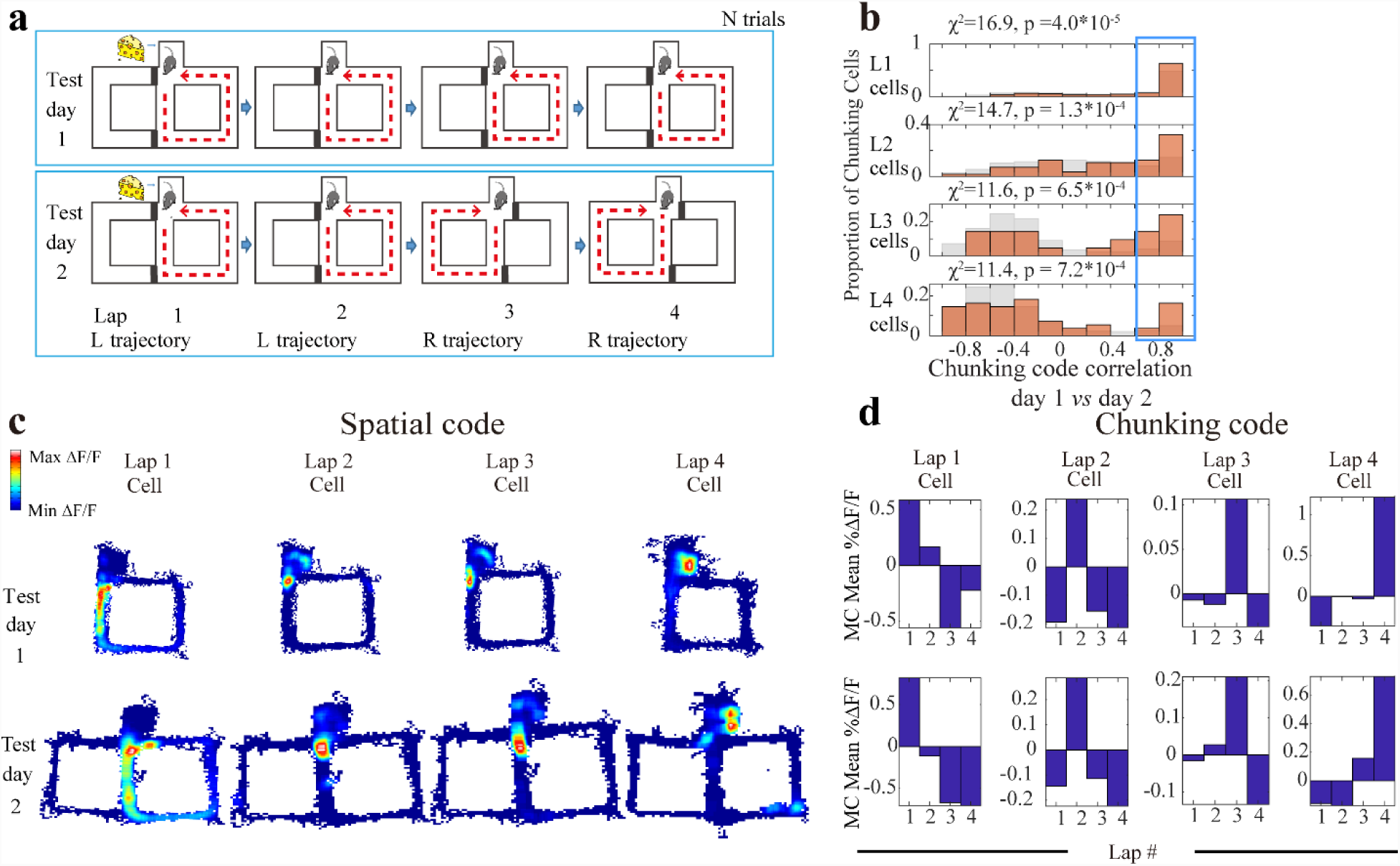
CA1 chunking cells tracks abstract “lap events” despite differential spatial trajectories. a) Alternation experiment: Day 1: standard 4-lap experiment and Day 2: alternating trajectory version. b) Chunking code correlations across the standard and alternating maze sessions (371 cells, n = 4 mice). See Fig. 1(l) for description and methods. c-d) Representative lap 1, 2, 3, and 4 neurons matched across standard and alternating maze sessions.

**Extended Data Fig. 8:**
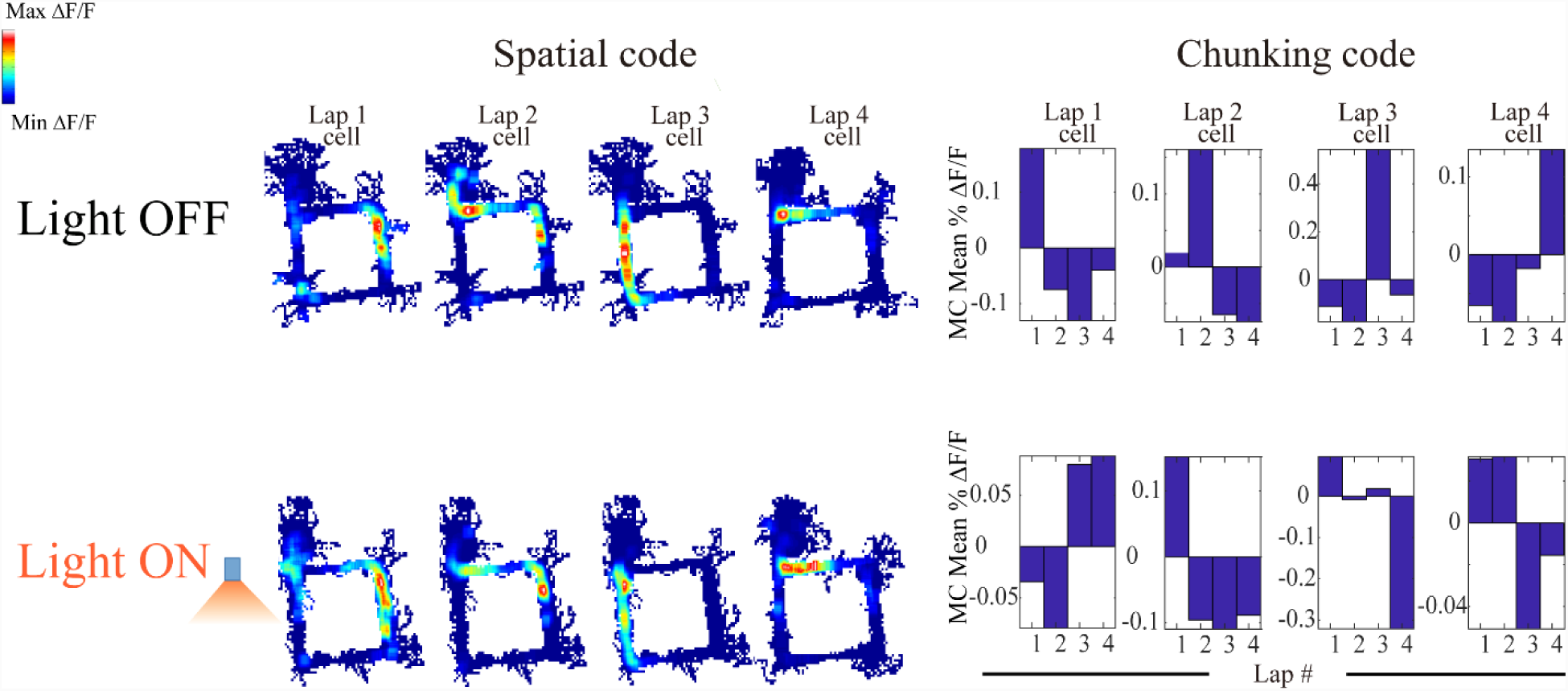
Chunking cells during MEC inactivation (more examples) Representative lap 1, 2, 3, and 4 neurons matched across Light-OFF and Light-ON trials.

**Extended Data Fig. 9:**
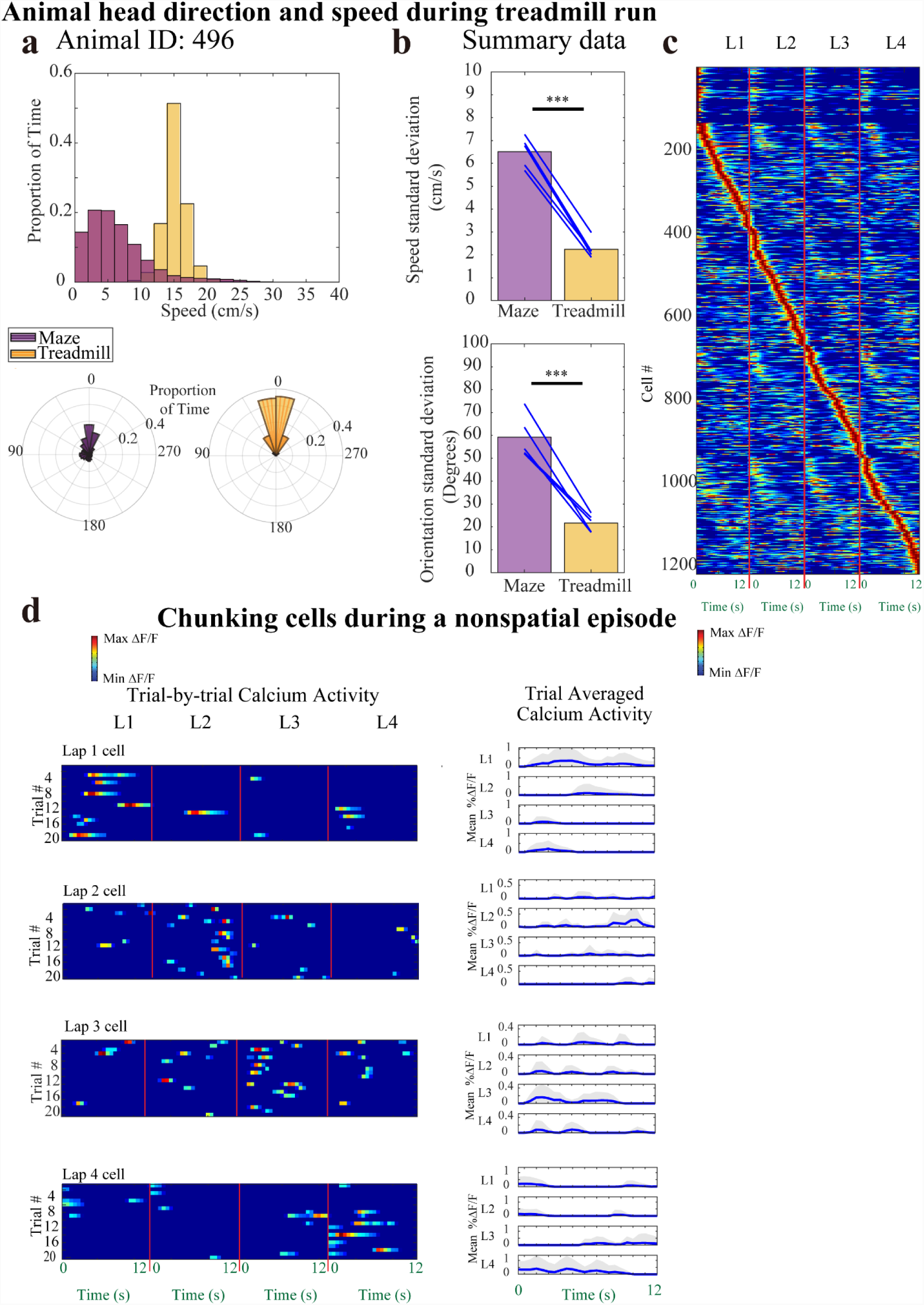
Chunking code during the treadmill period. a) Distribution of **top:** animal running speed, and **bottom:** animal head direction during the maze running portion (**purple**) versus the treadmill running portion (**yellow**) of the task, for animal 481 in 20 trials. b) Summary data: comparison of standard deviation of **top:** animal running speed (tstat = 19.65, df = 4, p = 4.0 * 10^−5^), and **bottom:** animal head direction (tstat = 9.32, df = 4, p = 7.4 * 10^−4^) during the maze running portion (**purple**) versus the treadmill running portion (**yellow**) of the task, for 5 animals. c) CA1 calcium activity sorted by spatial position and lap number, normalized and Gaussian smoothed (σ = 2s) calcium activity (1222 cells, n = 5 animals). d) Representative lap 1, 2, 3, and 4 neurons, temporally binned calcium activity during the treadmill period (0.5s bins). **Left** panel: trial-by-trial activity, **Right** trial averaged activity with mean ± SD. Standard deviation was cut off at 0 because negative activity does not exist.

**Extended Data Fig. 10:**
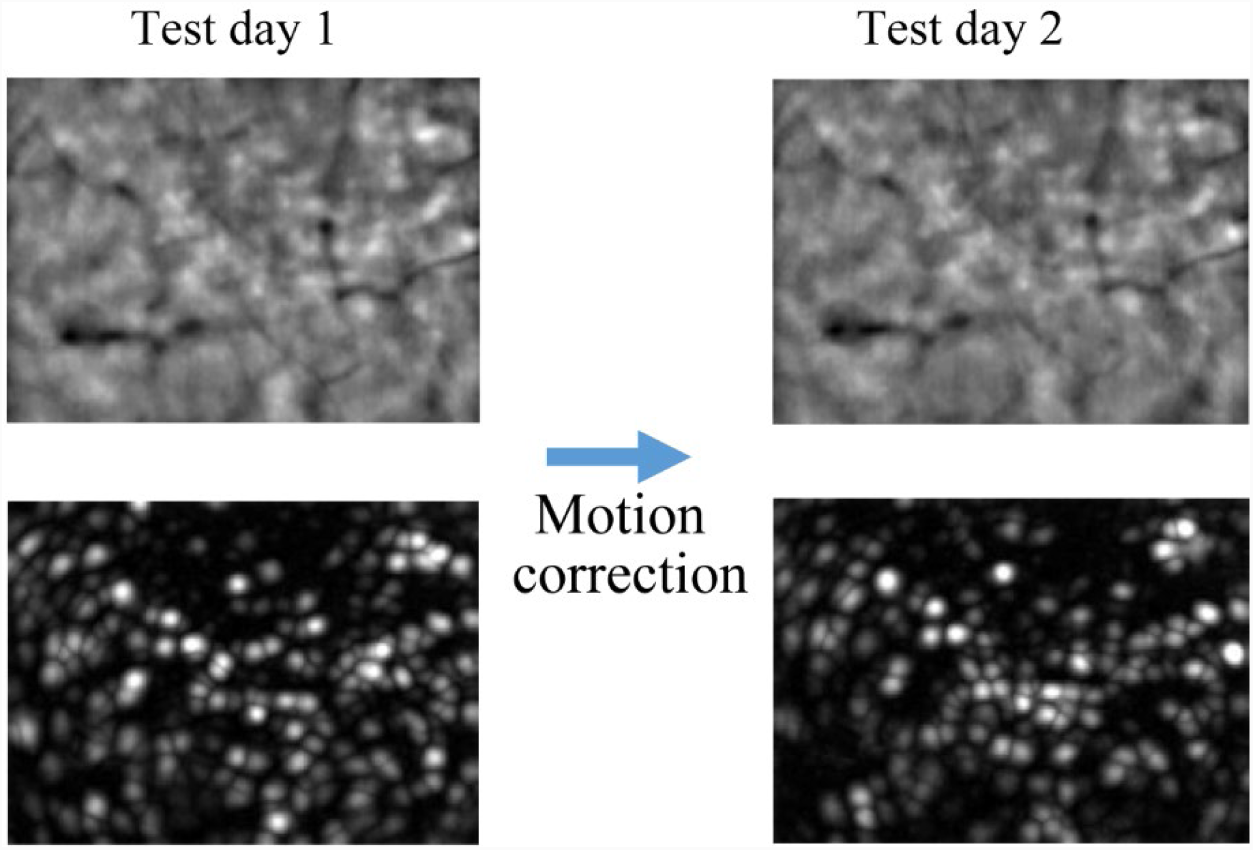
Method of image field of view registration across days.

## Materials and Methods

### Animals

All procedures relating to mouse care and treatment conformed to the institutional and NIH guidelines. Animals were individually housed in a 12 hour light (7pm-7am)/dark cycle. Nineteen male Wfs1-Cre mice aged between 2-4 months were food restricted to 85-90% normal body weight for the experiments. For each of the six main maze manipulation imaging experiments (fixed maze elongation experiment, random maze elongation experiment, lap addition experiment, spatial rotation experiment, treadmill experiment, spatial alternation experiment) the number of animals used (at least 4) is indicated in the main text for each experiment. In each of these experiments, at least two of these tested animals did not previously undergo any of the other manipulative experiments. The other animals were experienced animals from the other manipulative experiments. The exact number of animals used in each of these experiment is indicated in the text directly. Three pOxr1-Cre mice, aged 2-4 months, were also implanted with Inscopix microendoscope into CA1 for dual imaging and optogenetics experiments, and food restricted and trained in the same manner as the Wfs1-Cre mice.

### Histology and Immunohistochemistry

Mice were transcardially perfused with 4% paraformaldehyde (PFA) in phosphate buffered saline (PBS). Brains were then post-fixed with the same solution for 24 hours, and brains were sectioned by using a vibratome. Sections were stained by DAPI. Micrographs were obtained using a Zeiss AxioImager M2 confocal microscope using Zeiss ZEN (black edition) software.

### Preparation of Adeno-Associated Virus (**AAV**)

The AAV2/5-Syn-DIO-GCaMP6f was generated by and acquired from the University of Pennsylvania Vector Core, with a titer of 1.3*10^13 genome copy/ml. The AAV2/5-CamKII-GCaMP6f-WPRE-SV40 was generated by and acquired from the University of Pennsylvania Vector Core, with a titer of 2.3*10^13 genome copy/ml. The AAV2/2-EF1a-DIO-eNpHR3.0-mCherry was generated by and acquired from the University of North Carolina (Chapel Hill) Vector Core, with a titer of 5.3*10^12 genome copy/ml.

### Stereotaxic Surgery

Stereotactic viral injections and microendoscope implantations were all performed in accordance with Massachusetts Institute of Technology (MIT)’s Committee on Animal Care guidelines. Mice were anaesthetized using 500 mg/kg avertin. Viruses were injected by using a glass micropipette attached to a 10 µl Hamilton microsyringe through a microelectrode holder filled with mineral oil. A microsyringe pump and its controller were used to control the speed of the injection. The needle was slowly lowered to the target site and remained for 10 minutes after the injection.

For CA1 imaging experiments, unilateral viral delivery into the right CA1 of the Wfs1-Cre mice was aimed at coordinates relative to Bregma: AP: -2.0 mm, ML, +1.4 mm, DV, -1.55 mm. Wfs1-Cre mice were injected with 300 nl of AAV2/5-Syn-DIO-GCaMP6f. Approximately one month after injection, a microendoscope was implanted into the dorsal part of CA1 of the Wfs1-Cre mice aimed at coordinates relative to Bregma, at: AP: -2.0 mm, ML, +2.0 mm, and DV at approximately -1.0 mm. To implant at the correct depth, the cortex was vacuum-aspirated which resulted in the removal of corpus callosum, which is visible under surgery microscope as fibers running in the medial-lateral direction. The fibers of the alveus, which are visible as fibers running in the anterior-posterior direction, were left intact by the procedure.

For CA1 optogenetic and imaging experiments, 300 nL (AAV2/5-CamKII-GCaMP6f-WPRE-SV40) unilateral viral delivery into the right CA1 of the pOxr1-Cre mice was aimed at coordinates relative to Bregma: AP: -2.0 mm, ML, +1.4 mm, DV, -1.55 mm, and 300 nL (AAV2/2-EF1a-DIO-eNpHR3.0-mCherry) bilateral viral delivery into the MEC of these mice were aimed at coordinates relative to Bregma: AP: -4.85 mm, ML, ±3.45 mm, DV, -3.35 mm. Following these virus injections, the micro-endoscopy lens was implanted in the same manner for these dual optogenetic and imaging experiments, as described above for CA1 imaging experiments.

The baseplate for miniaturized microscope camera was attached above the implanted microendoscope in the mice. After experiments, animals were perfused, and *post hoc* analyses were examined to determine actual imaging position in CA1 (Fig. 1c, Fig. 3m).

### Apparatus description and Experimental conditions

The apparatus was a square maze 25 cm in length and width, with a 5cm wide track width, and 7 cm height. A 10 cm x 10 cm square reward box was located in one corner of the square maze. Sugar pellets (Bio-Serve, F5684) were placed in the reward box at the beginning of lap 1 of each trial. Four versions of this apparatus were used. Version 1 was used in Fig. 1-4 except Fig. 2. Version 2 used in Fig. 2 had a length elongation to twice the standard length (50 cm = 2 ×25 cm), but was otherwise identical to Version 1 in other dimensions. Version 3, used in Fig. 5’s treadmill experiment, had a 18 cm long treadmill installed in the arm of the maze that immediately faces the reward box (Fig. 5a). Version 3 otherwise used the same dimensions as Version 1. Version 4, used in Extended Data Fig. 7, had an 8-maze configuration, with the other square of the 8-maze being 25 cm in length and width as well. Version 4 otherwise used the same dimensions as Version 1. All maze experiments were done on under dim light conditions, with prominent visual cues within 50 cm on all sides of the box. Ca^2+^ imaging in the maze lasted at least 20 minutes in order to collect a sufficient number of Ca^2+^ transients to power our statistical analyses. The maze surface was cleaned between sessions with 70% ethanol. Immediately before and after imaging sessions, the mouse rested on a pedestal next to the maze.

The basic task in this manuscript is the standard 4-lap-per-trial task, where animals traversed round a square maze 25cm in length (1m journey in total) (Fig. 1d). The task was designed so that a sugar pellet reward was delivered manually to the reward box, at the beginning of lap 1, once every 4 such laps, which we call a single ‘trial’ (Fig. 1d). Identical motions were made on each lap, regardless whether a pellet was delivered or not. During the testing, animals completed 15-20 of such trials in repetitive succession without interruption. For any behavioral session in which the animal missed going into the reward box more than once in the entire sequence of runs (15 to 20 x 4 = 60 to 80 runs in total), the experiment was excluded. Crucially, for all experiments, animals first underwent task training before the final testing days. Training procedures are described below:

#### Habituation to reward in the maze

All behavior experiments took place during the animals’ dark cycle. All implanted mice were habituated to human experimenters as well as the experimental room for 1 week. At the same time, they were mildly food restricted and habituated to sugar pellet reward. The criterion for habituation to sugar pellets and the maze was running counter-clockwise around the maze and eating a sugar pellet in the reward port of the maze (described below) in 15 successive repetitions without missing a single pellet.

#### Reward periodicity-training

Animals were trained for approximately 8 days. If during any training day, the mice appeared unmotivated or too satiated to complete the 15 trials, that training day was repeated the following day. Animals were pre-trained for 2 days on the maze to habituate to receiving sugar pellet rewards in the reward port: on each of these days, they did a 1-lap-per-trial task, that is, they receiving reward every run around the maze, and ran 15 such trials. For the next 3 days (days 3-5), animals were trained to receive periodic rewards. On day 3, animals ran 15 trials of a 2-lap-per-trial task, that is, they receiving reward every 2 laps around the maze. On day 4, animals ran 15 trials of a 3-lap-per-trial task. On day 5, animals ran 15 trials of a 4-lap-per-trial task. Finally, animals ran 3 more days (days 6-8), 15 trials per day of a 4-lap-per-trial task, before they were considered well trained on the basic 4-lap-per-trial task.

#### Pre-versus Post-Training Experiment Protocol

In the particular case of the pre-versus post-trained animal experiment (Extended Data Fig. 4), animals that had only been habituated to the reward (described above) were immediately tested/imaged by running 15 trials of the standard 4-lap-per-trial task. Following this initial testing, these animals then underwent the reward periodicity-training (described above). Following periodicity-training, animals were tested/imaged again on 15 trials of the standard 4-lap-per-trial task, to compare the chunking cells seen post-training, compared to pre-training.

#### Reward on every lap experiment

Animals in this experiment were given a sugar pellet on every lap, completed a total of 60-80 laps total. This is equivalent to the total number of laps in the 15-20 trials of the 4 lap-per-trial experiment. This experiment did not require extra or task-specific training.

### TASK-SPECIFIC TRAINING

Each of the main maze manipulation experiments (fixed maze elongation experiment, random maze elongation experiment, lap addition experiment, spatial rotation experiment, treadmill experiment, spatial alternation experiment) required its own special ‘task’ training after completing the habituation and reward periodicity-training.

#### Fixed maze elongation experiment (Fig. 2)

For the fixed maze elongation experiments, animals were tested/imaged in a 2-day experiment. On Day 2, 2 hours prior to experimentation, animals were habituated (allowed to run) for 3 minutes on the distorted maze without any rewards.

#### Random maze elongation experiment (Fig. 2)

For the random maze elongation experiment, animals were tested/imaged on 28 4-lap trials. The maze was elongated on random laps of random trials, such that each of the 4 types of trials (SSSS, SSLL, LLSS, LLLL, where S denotes a “short” lap and L denotes a “long” lap) were presented in pseudorandom order (Extended Data Fig. 6 for full schedule) and appeared 7 times each within the 28 trials. Prior to test day, animals underwent 3 days (days (-3) to (-1)) of habituation training to the short and long laps, where SSSS, SSLL, LLSS, LLLL trials were presented randomly.

#### Five-lap-per-trial experiment (Fig. 3)

For the 5-lap-per-trial experiment, animals were tested/imaged in a 2-day experiment. These animals underwent 3 days (days (-3) to (-1)) of habituation training before the first test day. On the first 2 training days (day-3 to -2), animals each day ran 15 trials of a 5-lap-per-trial task. On the 3^rd^ day of training, (day (-1)) animals ran 15 trials of a 4-lap-per-trial task again, to get them habituated to test day.

#### Optogenetics experiment (Fig. 3)

Calcium imaging used the Inscopix nVoke miniature optoscope, occurring at 20 Hz. During periods of optogenetic manipulation as defined by our protocol (Fig. 3l), the Inscopix nVoke miniature optoscope’s orange light (590-650 nm) stimulation was turned on, at 10 mW/mm^2^ power, at a uniform and constant level. Orange light delivery was done manually and was turned on or off at the start of the relevant trial as soon as animals entered the box.

For optogenetic manipulation experiment, animals were tested/imaged in a single day. These animals underwent 2 days of habituation training before the first test day with 2 days in between each of the training days to allow recovery from the light. On each of the training days, animals each day ran 16 trials of a 4-lap-per-trial task with the light schedule according to the alternating schedule shown in Fig. 3l.

#### Spatial rotation experiment (Fig. 4)

For the spatial rotation experiment, animals were tested/imaged in a 2-day experiment. On Day 2, 2 hours prior to experimentation, animals were habituated (allowed to run) for 3 minutes on the distorted maze without any rewards.

#### Treadmill experiment (Fig. 5)

For the treadmill experiment, animals were tested/imaged in a single day. These animals underwent 6 days of habituation training running on the maze before the first test day. On the first day of training, animals ran 15 trials of a 1-lap-per-trial task. During each lap, the animal ran onto the first arm of the square maze, and ran for 12s (time period accurately indicated via Arduino) on the treadmill at a constant 14 cm/s, before running around the rest of the square maze and entering the reward box. On the next five days of training, animals ran 15 trials of a 4-lap-per-trial task again, with 12s on the treadmill, to get them habituated to test day.

#### Alternation maze experiment (Extended Data Fig. 7)

For the spatial alternation experiment, animals were tested/imaged in a 2-day experiment. These animals underwent 5 days (days (-5) to (-1)) of habituation training before the first test day. On the first 4 training days (day-5 to -2), animals each day ran 15 trials of a 4-lap-per-trial task where the laps alternated in their spatial trajectories according to Extended Data Fig. 7a. On the 5^th^ day of training, (day (-1)) animals underwent ran 15 trials of an ordinary (non-alternating) 4-lap-per-trial task again, to get them habituated to test day.

### Behavioral analysis and Ca^2+^ events detection

The animal’s position was captured by an infrared camera (Ordro infrared camcorder, 30 fps) via infrared light emitting diodes (LEDs) attached to the animal. Calcium events were captured at 20 Hz on an Inscopix miniature microscope. Imaging sessions were time stamped to the start of the behavioral recording by the turning on of an LED that is fixed to the animal, at the beginning of the session, and turning off of the LED at the end.

Analysis of the calcium images and extraction of independent neuronal traces were done akin to previous methods (*30, 41*). Specifically, the calcium movie was then binned 4x spatially along each dimension, and then processed by custom made code written in ImageJ (dividing each image, pixel by pixel, by a low-passed (r = 20 pixels) filtered version). It was then motion corrected in Inscopix Mosaic software 1.2.0 (correction type: translation and rotation; reference region with spatial mean (r = 20 pixels) subtracted, inverted, and spatial mean applied (r = 5 pixels)). A spatial mean filter was applied to it in Inscopix Mosaic (disk radius = 3), and a ΔF/F signal was calculated.

Four hundred (400) cell locations were selected from the resulting movie by PCA-ICA (600 output PCs, 400 ICs, 0.1 weight of temporal information in spatio-temporal ICA, 750 iterations maximum, 1E-5 fractional change to end iterations) in Inscopix Mosaic software. Region of interest (ROIs), half-max thresholded, that were not circular (if its length exceeded its width by > times) or smaller than 5 pixels in diameter (∼12 um), were discarded. For each ROI filter, pixels less than 75% of the filter’s maximum intensity were zeroed.

ΔF/F calcium traces were calculated for the resulting ROI filters for each processed movie. Slow variations in the calcium traces were eliminated by subtracting the median percentile ΔF/F value at each timepoint, this value calculated from the calcium trace values ±15s within this timepoint, similar to Ziv et al, 2013(*30*). The calcium trace was smoothed by 4-temporal bin rolling average (each bin 50 ms). Significant calcium transients (Fig. 1c) were detected as traces that exceeded 3 Standard Deviations above baseline, and furthermore, remained above 1.5 Standard Deviations above baseline for at least 500 ms. The rest of the ΔF/F calcium traced, aside from its significant transients, were zeroed similar to Dombeck, 2010. Only cells that had a total of at least 25 significant transients during the entire session and nonzero activity in at least 10 trials separately were considered for further analysis in this study. In the sole case of the treadmill experiment, a lesser total of at least 10 significant transients was used, since the cumulation of all the treadmill periods was only 12-16 min (15-20 trials).

### Chunking cell calculation

#### A) Calcium event filtering

For each CA1 cell detected, the calcium activity was filtered so that only activity occurring while the mice were in an active state (animal speed > 4 cm/s) were analyzed further. The behaviorally tracked times of interest were also filtered in this way, considering only the times with animal speed > 4 cm/s. The maze was divided into 9 spatial bins: the reward box (spatial bin of length and width 10 cm) was one spatial bin, and each of the 4 arm lengths of the maze were divided in half (8 spatial bins, each of which was 12.5 cm in length and 5 cm in width).

Next, for each identified cell, individual calcium activity epochs were analyzed by calculating the mean calcium activity in each of the 9 spatial bins during each individual lap across trials. Thus, for a session of 15-20 trials, there were 540 to 720 calcium activity epochs in total (15 to 20 × 9 ×4 = 540 to 720).

Each CA1 neuron possesses a spatially tuning, and in this model, the spatial tuning was captured by a parameter **p** defined as the probability of having nonzero calcium activity in each separate spatial bin. **p** was calculated for each neuron for each of its spatial bins. It differed for different spatial bins, reflecting the spatial code.

#### Linear model fitting

For each activity epoch for each neuron, the mean ΔF/F calcium activity, the mean speed (s), and the head direction tuning (o) were calculated. The nonzero calcium activity epochs were fit by a linear regression of the mean ΔF/F calcium activity versus speed and head direction tuning. In this regression, the coefficients *a, b, c* were fit:

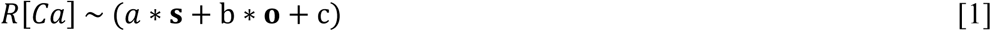

Where R[Ca] is the mean ΔF/F calcium activity level of this neuron during this activity epoch. **s** is the mean speed of the animal during this activity epoch, and **o** is the head orientation deviation from the preferred head orientation of this neuron during this activity epoch. In Matlab code, we used the function: `fitrlinear` with lambda = 0.01, to fit the equation [1] using regularized linear regression applied to the calcium activity epochs of all cells.

#### B) Identification of Chunking cells

For each identified cell, we shuffled its calcium transients across the lap epochs, such that the probability of assigning any particular calcium transient into any particular lap epoch varied according to equation [1]. Calcium transients were only shuffled (using randperm in matlab) between different epochs taking place in the same spatial field in order to preserve **p**. We checked that this shuffle generation procedure gave a mean ΔF/F calcium activity level that matched the model-predicted (equation [1]) calcium activity level (Extended Data Fig. 3c). These shuffles simulated the calcium activity of the cell explained by spatial field (**p**), head direction (**o)** and animal speed (**s**). A total of 5000 such shuffles were computed, and a ‘model-explained mean ΔF/F calcium activity level’ was computed as

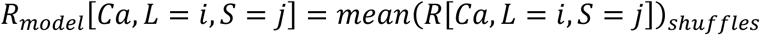

Where *R*[*Ca, R* = *i, S* = *j*] is the ‘model-explained calcium activity’ computed as the mean activity in lap *i* and spatial bin *j* across all the shuffles for this cell.

For every neuron on all four individual laps, the model-explained mean calcium activity level in each individual spatial bin was subtracted from the real mean ΔF/F calcium activity, to yield ‘model corrected’ (MC) ΔF/F calcium activity which excluded spatial, mean speed, and mean head direction tuning (Fig. 1h). Thus, this model corrected effect would mainly reflect difference in calcium activity due to lap number.

For every neuron, the model-explained mean ΔF/F calcium activity level was subtracted from the mean ΔF/F calcium activity level obtained from the 5000 shuffles, to yield a <u>distribution</u> of MC ΔF/F activities for chance level statistics. Cells whose peak, lap-specific MC ΔF/F was outside the 95^th^ percent confidence of shuffled MC ΔF/F were called ‘**significant chunking cells**’.

If the peak MC calcium activity happened to occur during the reward eating lap (lap 1) while the animal was in the reward box spatial bin, then the peak MC calcium activity from the next highest spatial bin was selected, because we excluded cell activity that was directly driven by reward eating.

We also considered looking at raw ΔF/F activity of these chunking cells to examine if similar results were obtained (Extended Data Fig. 5).

### Spatial information

The tracked positions were sorted into 16 spatial bins of size 6.25cm x 5cm around the track and 4 spatial bins of size 5cm x 5cm in the reward box and the mean ΔF/F calcium activity of each CA1 cell was determined for each bin. The bins which had animal occupancy < 100 ms were considered unreliable and discarded from further analysis. Without smoothing, the spatial tuning was calculated for each cell according to:

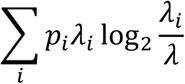

Where *λ*_*i*_ is the mean ΔF/F calcium activity of a unit in the *i*-th bin, *λ* is the overall ΔF/F calcium activity, and *pi* is the probability of the animal occupying the *i*-th bin for all *i.* This formulation, derived in Skaggs et al, 1993(*42*), was applied to calcium activity levels, which have a known monotonic relationship to spike rates (Chen et al, 2013). All cells’ event times were shuffled 2000 times in an analogous manner to Wills et al, 2010 by shifting the calcium activity time series around the position data by a random translation of > 20 s and less than the session duration minus 20s. Cells with significant spatial information were determined above the 95^th^ percentile of all shuffles.

### Registering cells across days

Our approach to register cells across days was to do so on the basis of the anatomy of the field of view seen on both days (i.e. the pattern of blood vessels, etc.), rather than on the spatial locations of cells directly (Extended Data Fig. 10). Then, after an appropriate image registration was found for the fields of view based on anatomy, the ROIs on day 1 were identified, and calcium traces were calculated based on the resulting ROI filters for day 1 applied directly to the processed movie on day 2. To register two movies across days, a mean projection of the ImageJ filtered and motion corrected movie (see above methods) on each day was computed, and these two movies were registered with respect to one another by the Inscopix Mosaic motion correction software.

### Chunking and Spatial correlations across days

For chunking correlations across days: for a given significant chunking cell on day 1, its chunking code (defined in the main text) was concatenated into a vector. A similar vector was produced for this same cell on day 2. This was done for each significant chunking cell matched across days. The Pearson correlation between the day 1 chunking code vector and day 2 chunking code vector was calculated to examine chunking code preservation across days. The day 2 chunking code vector was produced from the same spatial bin as day 1 to allow for direct chunking code comparison, except for the spatial rotation and spatial trajectory alternation experiment. In these cases the spatial bin in which peak activity occurred were calculated anew, since the space was substantially changed in these experiments relative to room cues.

For spatial correlations across days: The raw calcium events, speed filtered (> 4 cm/s) were sorted into the 9 spatial bins defined above and the calcium activity level of each neuron was determined for each bin, and an activity map composed of all the spatial bins was produced. The activity maps for each individual chunking cell was treated as a vector (list of numbers) and Pearson correlation between the spatial activity maps of the two days was calculated.

### Statistics

All statistical tests in this study were two tailed.

